# GridFree: A Python Package of Image Analysis for Interactive Grain Counting and Measuring

**DOI:** 10.1101/2020.07.31.231662

**Authors:** Yang Hu, Zhiwu Zhang

## Abstract

Grain characteristics, including kernel length, kernel width, and thousand kernel weight, are critical component traits for grain yield. Manual measurements and counting are expensive, forming the bottleneck for dissecting the genetic architecture of these traits toward ultimate yield improvement. High-throughput phenotyping methods have been developed by analyzing images of kernels. However, segmenting kernels from the image background and noise artifacts or from other kernels positioned in close proximity remain challenges. In this study, we developed a software package, named GridFree, to overcome these challenges. GridFree uses an unsupervised machine learning approach, K-Means, to segment kernels from the background by using principal component analysis on both raw image channels and their color indices. GridFree incorporates users’ experiences as a dynamic criterion to set thresholds for a divide-and-combine strategy that effectively segments adjacent kernels. When adjacent multiple kernels are incorrectly segmented as a single object, they form an outlier on the distribution plot of kernel area, length, and width. GridFree uses the dynamic threshold settings for splitting and merging. In addition to counting, GridFree measures kernel length, width, and area with the option of scaling with a reference object. Evaluations against existing software programs demonstrated that GridFree had the smallest error on counting seeds for multiple crops, including alfalfa, canola, lentil, wheat, chickpea, and soybean. GridFree was implemented in Python with a friendly graphical user interface to allow users to easily visualize the outcomes and make decisions, which ultimately eliminates time-consuming and repetitive manual labor. GridFree is freely available at the GridFree website (https://zzlab.net/GridFree).

## Introduction

Grains are the staple food for more than four billion people worldwide. As population increases each year, so does the need for higher grain yields. Investigating the genomic factors controlling grain yield has become the principal methods of breeding programs. Previous studies indicate that grain yields are positively correlated with particular grain traits related to kernel characteristics, such as kernel length, width, area, and weight. Larger kernels contain more nutrients to feed the embryo before the plant establishes a root system and leaves to support itself (Trinetta 2016). Kernel size correlates with various seed functional traits, such as seminal root length and total root weight, and larger seeds exhibit improved seedling establishment and shoot weight (Thomas *et al.* 2016). Kernel size also affects germination capacity, offspring seedling establishment, and seedling germination times (Zohaib *et al.* 2018). In general, kernel size positively correlates with kernel weight (Abdipour *et al.* 2016).

Kernel size and shape also impact end-use quality, such as milling yield, protein content, and consumer preference (Gupta *et al.* 2006). In many markets, large and plump grains are more desired. Theoretical models predict that milling yield could increase when spherical grain shapes and large sizes are optimized (Gegas *et al.* 2010). Kernel shapes also create market classes for some grains. For example, Southeast Asian and South Chinese consumers prefer long-grain rice, but Japanese, Korean, North Chinese, and Taiwanese consumers prefer short-grain rice (Suwannaporn and Linnemann 2008).

Quantitative Trait Locus (QTL) analysis can reveal correlations between grain traits and polymorphisms of DNA. Grain breeders apply QTL analysis to identify and sequence the actual genes that cause grain kernel size variation(Gegas *et al.* 2010; Liu *et al.* 2016; Sun *et al.* 2019). A challenge in implementing QTL analysis is evaluating the phenotypic traits. Accurate measurements of grain kernel sizes are critical for QTL analysis. Many studies manually measure grain samples (Redona and Mackill 1998; Chen *et al.* 2016, 2019; Li *et al.* 2019), with sample sizes ranging from 20 to 50 per variety. However, evaluating kernel traits by manual measurements presents two distinct disadvantages. First, the process is laborious and tedious. Second, only small sample sizes are manageable and may not reflect the complete variation in kernel size of the variety of interest.

Using high-throughput phenotyping technology to evaluate grain kernel traits was first introduced in the late 1990s (Campbell *et al.* 1999). Since then, the cost of high-throughput technology has dramatically decreased, allowing for a more comprehensive range of applications in plant research (Yang *et al.* 2013). Current high-throughput methods include devices that generate digital images of grain samples and image processing software programs or tools that estimate grain kernel traits, saving breeders time and labor compared with manual methods. High-throughput methods are capable of handling large sample sizes such as 1000 kernels (Yin *et al.* 2015).

Devices for obtaining digital images of grain kernels include scanners (Gegas *et al.* 2010; Liu *et al.* 2016; Desiderio *et al.* 2019; Ma *et al.* 2019; Sun *et al.* 2019), cameras (Campbell *et al.* 1999), and other photographic apparatus (Yin *et al.* 2015; Cabral *et al.* 2018). The next step, processing grain kernel images, is a critical part of the high-throughput method. However, two significant challenges remain. The first challenge is accurately separating objects of interest (kernels) from other parts of the image, including shadows and contaminations such as harvest rubble. The second challenge is correctly separating adjacent objects of interest. Researchers spend great amounts of effort to manually spread grains apart prior to using a high-throughput imaging method to obtain accurate measurements of traits, such as 1000-kernel weight for a QTL study (Yin *et al.* 2015).

Some researchers use either computer software programs or smartphone apps that count kernel numbers and estimate grain kernel traits in given digital images (Rasband 1997; Tanabata *et al.* 2012; Whan *et al.* 2014; Komyshev *et al.* 2017; Gao *et al.* 2018; Wu *et al.* 2018). Most of these software packages have strict requirements for image background and grain distribution. SmartGrain (Tanabata *et al.* 2012), a phenotyping software program, can estimate kernel sizes; however, obtaining accurate results requires the labor-intensive task of spreading grain kernels evenly. Additionally, SmartGrain can only process images with a dark background. In contrast, SeedCounter, an Android App that counts the number of grain kernels from images, can only process images with a light background (Komyshev *et al.* 2017). SeedCounter uses a contour method to segment kernels. The contour method identifies boundaries of areas that consecutively have the same pixel value in an image. However, the method cannot separate adjacent grain kernels. GainTKW (Wu *et al.* 2018) is a system that combines an Android app and an electrical scalar. The scalar measures the weights of kernels; the app takes kernel photos and then processes the digital images. The app uses a Lab-Otsu color-index approach (Otsu 1979) to convert regular red-green-blue (RGB) images into binary images. Then, the app uses K-Means method to classify the image into two clusters to separate kernels from the background. The marker-controlled watershed method and normal distribution are applied as thresholds to segment adjacent grain kernels. Wu *et al.* (2018) showed that the app could process images with dark backgrounds. However, for light-background images, the app segmented the background into blocks and counted them as objects.

Besides, there are also computer platforms developed for extracting phenotypic features from other plants’ digital images. Miller *et al.* (2017) developed a web-based application that extracts size features from maize images, such as size estimation of maize ear and cob. The application adopts principal component analysis (PCA) on extracted sizes of maize. Klukas *et al.* (2014) reported an open-sourced platform developed in Java named Integrated Analysis Platform (IAP). The system requires an image without plants as reference background, then separates plants using the reference background. Outputs include estimated phenotypic features, such as height, leaf count, top/side areas, and so on. Root growth is an important phenotypic feature to monitor the growth of plants. French *et al.* (2009) published a software tool named RootTrace that estimate plant root length. The software applies a pruned graph that traces plant roots from user-defined start points to the root tips.

Several software packages are tolerant of both dark and light backgrounds but have weaknesses when processing images with adjacent objects. GrainScan is a software program that requires users to input estimates of the upper and lower limits for the kernel area as references to process adjacent grains (Whan *et al.* 2014). Museed (Gao *et al.* 2018) is an Android App that applies K-Means to cluster grain kernels and a marker-controlled watershed method, which uses Euclidean distance between area boundary and area center, to segment adjacent grain kernels. The app uses normal distributions as thresholds to determine the trajectory of segmentation lines between adjacent grain kernels. Another software program, ImageJ (Rasband 1997), works in a variety of backgrounds and can count and estimate kernel sizes. But ImageJ requires that grain kernels are spread evenly and requires users to choose color-index approaches to segment kernels from the background (Mussadiq *et al.* 2015).

In this study, our first objective was to develop methods for (a) selecting pixels corresponding to objects of interest without explicitly examining a variety of color-indices, and (b) conducting segmentation without the restriction of normality of segmentation. Our second objective was to implement these methods in the development of a user-friendly software package with a graphical user interface (GUI). The GUI would allow users to select pixels of interest and visually perform interactive segmentation by checking boxes and sliding bars with real-time preview. With our software package, named GridFree, researchers can choose any background that contrasts with their objects of interest. Adjacent seeds are acceptable as long as none are overlapping. With GridFree, users can also compare different images by using a reference object. GridFree is an open-sourced package with a MIT license. All supporting documents can be found at the GridFree website (https://zzlab.net/GridFree), including user manual, tutorial, demonstration images, and source code.

## Results

Herein, we explain the design, the implementation of GridFree, and how the program works. Following, we present the results of our tests examining GridFree’s effectiveness in counting grain kernels and measuring kernel traits. We compare GridFree’s counting error rates with existing software programs. We also present the applications of segmenting and measuring other objects, including E Coli colonies, blood cells, spores, and field plots.

### Extraction of pixels of interest

A variety of vegetation indices have been developed and presented in the literature to extract pixels of interest (**Table S1**). Some of these indices can be better measurements of image content than the raw channels for specific purposes. We classify these color indices by the channels (RGB) and properties (**Table S2**). The properties were grouped into ten categories, including a ratio of one channel over another channel, and a ratio of one channel over the sum of others. The three channels and the ten properties formed a total of 30 color indices. Some of these indices were well investigated. For example, the proportion of green over the total (Green + Red + Blue), one of Proportion of Total (POT), was defined as Greenness (Wan *et al.* 2018).

Some of these color indices are more correlated than others (**Figure S1**). Using the three channels simulated indecently, we found Proportion Among Two Bands (PAT) is more correlated to the Average Difference Proportion (ADP). Based on the relationship among the 30 color indices, we selected 12 indices covering three raw channels and four properties (**Figure S2**), including PAT, Difference (DIF), Ratio Over Other (ROO), and Golden Ratio (GLD). The images of principal components (PCs) derived from the 12 indices were almost identical to the PC’s images derived from the 30 indices (**Figure S3**).

GridFree conducts Principal Component Analysis (PCA) on both the 12-color index and the three RGB channels separately. The values of PCs are scaled to 0 to 255 for display as 8bit gray images. The 12-color index PCs are displayed as thumbnail array on the control panel with four rows and three columns. Each PC image displays on the display panel once users click the corresponding thumbnail. The RGB PCs are displayed differently. The first two RGB PCs can be displayed by moving the sliding bar to the left end for PC1, or to the right end for PC2. PC3 was ignored as there is barely information left outside the first two PCs for the three RGB channels. When the sliding bar is at the center, the display panel demonstrates the selected index PC (**Figure 1a** to **1h**).

**Figure 1.**
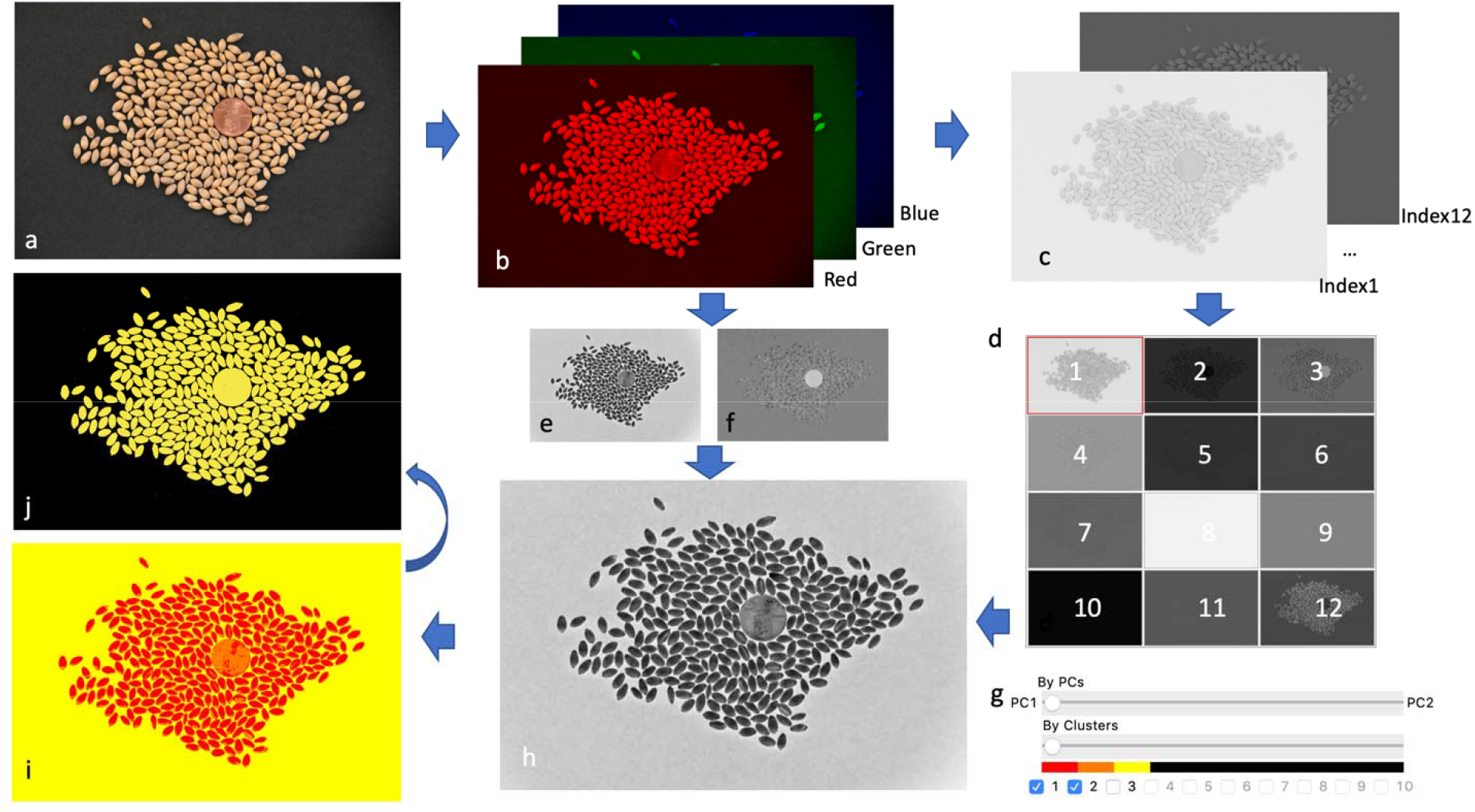
Selection on pixels of interested using principal component analyses and K-Means. GridFree inputs are RGB images (a). GridFree automatically separates the three RGB channel (b), computes 12 color-indices and implement PCA on the three RGB channels and the 12 color-indices separately (c). GridFree convert pixels PC values into gray image (ranges 0 to 255) The 12 color indices PC images are displayed as thumbnail buttons for selection (d). PC1 (e) and PC2 (f) of RGB channels can be displayed using the sliding bar (g). RGB PCs can be combined with the 12 color-indices PCs using the weight controlled by the sliding bar. The combined PC is instantly displayed (h). Cluster analysis (i) is controlled by sliding bar (g) for number of K-Means clusters. The final selection of pixels of interest (j) is controlled by checkboxes (g).

GridFree allows users to consider both RGB and index PCs. For a selected index PC, the sliding bar’s default position is at the center, indicating both RGB PC1 and PC2 weight zero. When the sliding bar is moved to the left, the weight on RGB PC1 increases and the weight on the selected index PC decreases. The weight on RGB PC2 remains at zero. When the sliding bar is to the left end, RGB PC1 is weighted 100%, and the selected index PC is weighted 0%. Similarly, when the sliding bar is moved to the right, the weight on RGB PC2 increases and the weight on the selected index PC decreases. The weight on RGB PC1 remains at zero. When the sliding bar is to the right end, RGB PC2 is weighted 100%, and the selected index PC is weighted 0% (**Figure 1g to 1h**).

For every clicking on an index PC thumbnail, or positioning the RGB PC sliding bar, the display panel instantly demonstrates the selection of Pixels Of Interest (POI) to serve as the first level of filtering. The second level of filtering is to conduct K-Means cluster analysis (**Figure 1d**), allowing users to choose the clusters to define the POI (**Figure 1e**). All filtering processes can be conducted on GridFree’s user-friendly GUI. The user interface consists of three panels: the display panel on left, the filter panel at right, and the control panel at the bottom. The display panel shows the raw image after loading by clicking the Image button in the control panel. The raw image can be switched to other filtering images and outcome images by clicking the corresponding buttons, including PCs, Clusters, Selected, and Outcome. The filtering of POI is operated at the top of the filter panel.

The default K value is one. The display panel instantly shows the results (**Figure 1i**) of changing K values with the cluster sliding bar. Checkboxes are used for selecting clusters. The first cluster is checked, and the other clusters are unchecked by default. The display panel instantly shows the selected clusters’ results (**Figure 1j**). The selected clusters are displayed in yellow; the other clusters are displayed in black.

### Segmentation

After users complete the filtering on PCs and clusters and click the Segment button, the segmentation results are displayed in two places, the display panel and the bottom of the filter panel (**Figure 2**). The display panel shows the segmentation as boxes outlined in red. The filter panel displays the scatter plot of the segments as their total POI against diagonal size. A segment can be selected by either clicking the segment on the display panel or clicking the dot on the filter panel. A segment is highlighted in both places simultaneously when the segment is selected on either place. Segmentation consists of three levels of filtering. The first level sets a reference for converting pixels to millimeters. The physical area of the reference (default = 255 mm^2^) is used to determine the actual size of a pixel. When the reference segment is selected and the reference box is checked, the segment is labeled as “Ref” after clicking the Segment button. To remove the reference, uncheck the “Ref” box, and click the Segment button.

**Figure 2.**
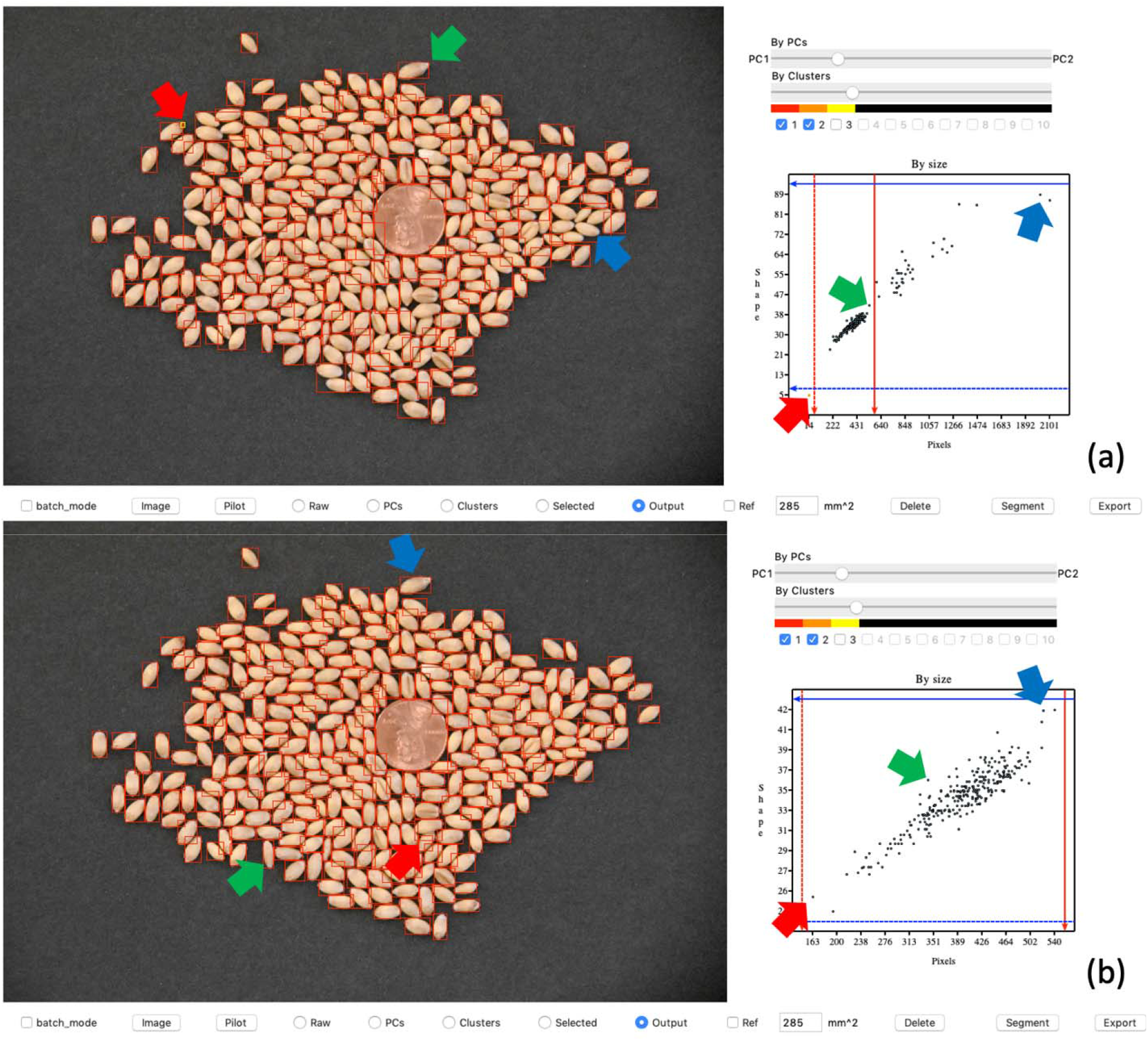
Interactive segmentation demonstration on GridFree’s graphical user interface. The left side of the user interface displays the image outputs of segmentations. The top image (a) is the result of an initial default segmentation. The blue and green arrows point to the greatest diagonal size of adjacent wheat seeds and that of single wheat seeds, respectively. The red arrow indicates the wheat seed with the smallest area. In scatter plot (a), users move the red and blue bars, through a click-and-drag motion with the mouse, to the locations where blue and green arrows point to set thresholds for further segmentation. The bottom image (b) is the result of the segmentation after setting the threshold guidance. Plot (b) displays a good relationship between the wheat seed areas and their diagonal sizes. The blue, green, and red arrows point to the biggest-, medium-, and smallest-sized wheat seeds in the segmentation output. The adjacent kernels in the display image (a) have been segmented individually in display image (b).

The second level of segmentation filtering is deletion. Multiple segments can be selected by clicking while holding the Shift key. These selected segments can be deleted by clicking the Del button. The deletion function is helpful when selected POI include both POI and non-POI that cannot be separated. The delete operation should be used with caution because, currently, GridFree has no option for restoring the deleted segments without redoing the entire segmentation process.

The third level of segmentation filtering is adjusting the size of the segments to match the actual grain kernels. We demonstrate the initial segmentation (**Figure 2a**) and the customized segmentation (**Figure 2b**). The horizontal axis represents the POI area of segments; the vertical axis represents the size of segments as the length of their diagonal. Both area and size are calculated as the number of pixels. Users can click on any of the dots in the preliminary segmentation plot to examine the accuracy of segmentation in the display image. Outliers can be easily identified by examining the data points (dots) that lie either above or below the central cluster of data points. In this demonstration, data points that lie above the central cluster include comparatively larger segments containing multiple adjacent grain kernels that require further segmentation. In contrast, data points that include comparatively smaller segments lie below the central cluster. These smaller segments are either the background noise artifacts that need to be removed or segments that need to be combined to create larger segments.

Users can set upper and lower boundaries for both area and size for segmentation. In the scatter plots, area boundaries are indicated by the red lines and size boundaries by the blue lines. Users can change the boundaries by clicking-and-dragging. After clicking the Segment button, segments larger than the thresholds (solid lines) are further divided and segments smaller than the thresholds (dash lines) are joined or deleted if the combined segments fail to satisfy the thresholds. Setting thresholds and clicking the Segment Button can be performed repeatedly until the user is satisfied with the outcome.

Once the final segmentation is generated, the user clicks the Export button to save the final output as both images and a Microsoft Excel file formatted as a comma-separated values (CSV) spreadsheet file. The Excel CSV output file contains the details of kernel (or item of interest) traits that can then be organized into spreadsheet cells and columns for analyses and data transfer. The output information includes two types: (1) all segmented kernels labeled by number and their associated lengths, widths, and areas in the number of pixels and millimeters (mm are computed only if users assigned an object as a reference); and (2) the summation, mean values, and standard deviations for color-index results and RGB channel separation.

### Comparing GridFree with existing software programs for counting seeds

To evaluate and validate GridFree’s effectiveness in segmenting adjacent grain kernels, we examined images with four types of grain kernels—black bean, canola, chickpea, and lentil. The numbers of grain kernels were 100, 199, 490, and 1001 for these species, respectively. Each grain kernel image contained two types of spatial patterns—adjacent and distinctly separate. Two images had a light or white background; the other two had a black background. Black beans and canola seeds are black kernels and were placed on white backgrounds. Chickpea and lentils are brown kernels and were placed on black backgrounds.

We also conducted a counting experiment on the grain kernel images to compare GridFree with one application on a smartphone (SeedCounter) and three computer software programs (SmartGrain, GrainScan, and ImageJ). We downloaded SeedCounter to an Android OS smartphone (OnePlue 7 Pro). We firstly use SeedCounter to count the number of seeds for each kernel scatter pattern setting (black beans and canola seeds). In addition to the existing background (in this case, it is a piece of white paper) SeedCounter requires the image to contain another background with different color as the white paper’s background. Once SeedCounter successfully delivered the counting result, we then use the same device to take pictures for other software programs. We cannot access the photo taken by SeedCounter, only be able to screenshot its output results. Therefore, we need to retake pictures for every seed scatter setting for other applications. For each case processed by SeedCounter, we attuned the calibration to make sure the application can process the image successfully. We selected the watershed algorithm in SeedCounter for image processing, and the paper is set to A4. We downloaded, installed, and ran SmartGrain and GrainScan on a Windows OS laptop to count the images’ grain kernels. We used the default thresholding for item sizes in GrainScan and SmartGrain. For ImageJ, we ran the evaluation on a Mac OS laptop. We used Otsu thresholding (Otsu 1979) in ImageJ for image pre-processing, the watershed algorithm to process the corresponding binary image, and set 75-infinity for the object size threshold.

We found that GridFree, GrainScan, and ImageJ can identify grain kernels on both light and dark backgrounds. SeedCounter can only recognize items on a light or white background; SmartGrain can only detect grain kernels on a dark or black background (**Figure 3**). GridFree exhibited the highest counting accuracies when compared with the four other methods. Among the two white background images accepted by SeedCounter, the counting errors for the black beans were less than the counting errors for the canola seeds. GainScan performed better on dark backgrounds than light backgrounds. SmartGrain showed higher accuracy counting spread-out grain kernels than adjacent grain kernels, which agrees with the (Tanabata *et al.* 2012) study. We observed that ImageJ exhibited an overwhelming number of errors when counting grain kernels on the chickpea image, likely due to the light reflected off the image surface.

**Figure 3.**
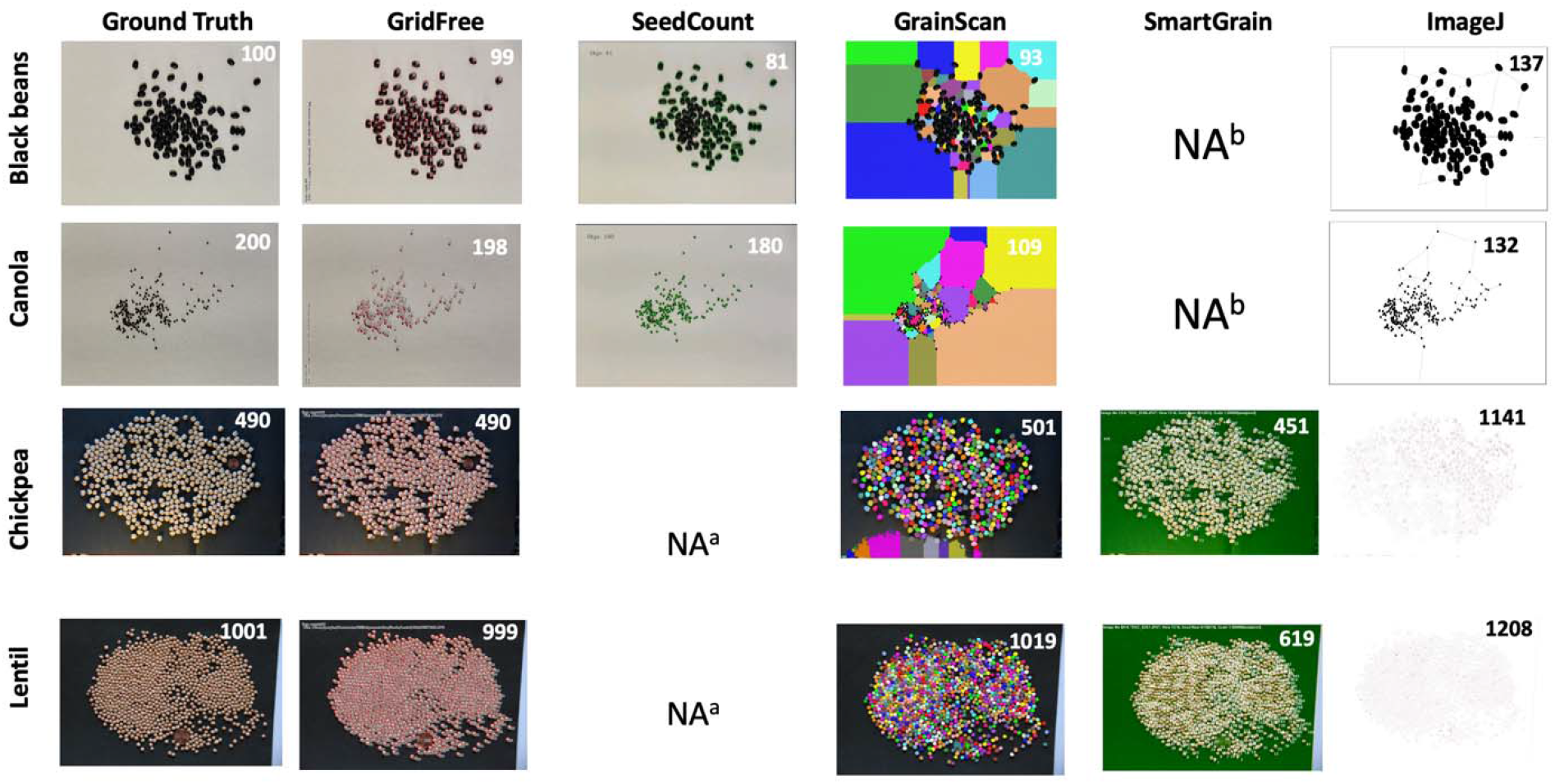
Comparison of counting results on seed images for five software. Comparison of GridFree’s counting accuracy with four other software packages, based on images of four different types of seed kernels: black bean, canola, chickpea, and lentil. ^a^The package does not accept black backgrounds ^b^The package does not accept white backgrounds

We replicated the experiment with multiple images of alfalfa seeds on a white background and made comparisons among GridFree, GrainScan, ImageJ, and SeedCounter. We conduct the same process as we did for the four types of grain kernels that use SeedCounter to process the seed scatter pattern. Once it successfully outputs counting results, we use the same device to take images for other applications. The number of seeds was set as four levels: 100, 200, 500, and 1000. The seeds for each level were randomly positioned ten times, and images were taken accordingly. Every image contained both spread-out and adjacent alfalfa seeds. Results showed that GridFree exhibited the least amount of counting errors across the four seed-count levels compared with the other three methods (**Figure 4**).

**Figure 4.**
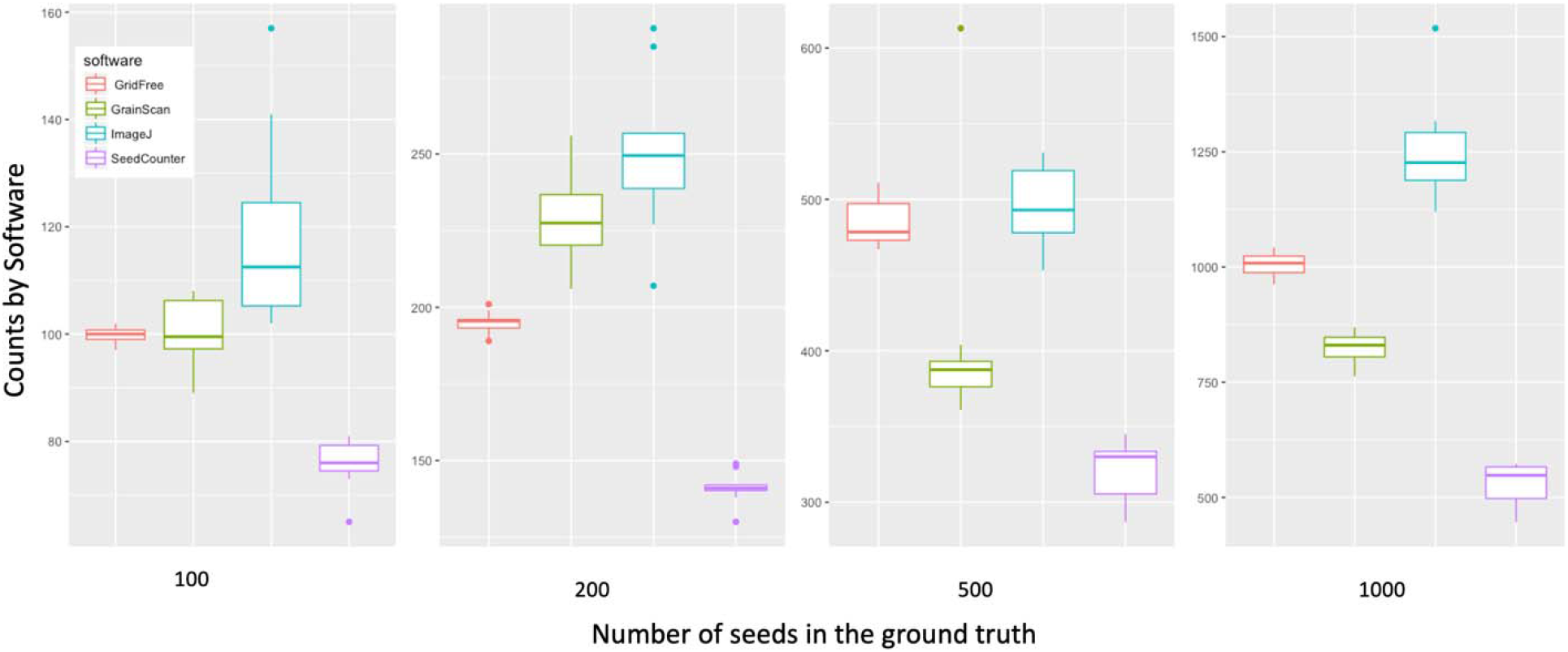
Comparison of counting accuracy on alfalfa seed images for four software programs/applications. Comparison of GridFree’s counting results with three other application, using images with different levels of seed counts. A total of 40 alfalfa seed images varied by number of seeds, with 10 images each of 100, 200, 500, and 1000 seeds.

### Seed measurements

GridFree produces measurements on objects, including total area, length, and width. These measurements use pixels as the units. When a reference is used, GridFree also outputs these measurements in the actual physical unit specified. The length is measured as the longest distance between two points at the edge of an object. The width is measure as the longest line between two points on the edges that are perpendicular to the length. The relationships among these measurements are displayed (**Figure S5**) for the wheat kernels used in the demonstration (**Figure 1**). The distributions of the three measurements are displayed on the diagonal, with units in mm for length and width and mm^2^ for the area. The correlations among area, length, and width are presented as scatter plots in the lower triangle; their Pearson correlation coefficients are displayed in the upper triangle.

We validated the size measurement accuracy of GridFree using manually measured wheat kernels. Wheat kernels are placed on a black paper, and an Android phone, OnePlus 7 Pro, took the image. The camera was set as taking a regular picture. The image is presented in Figure 5, the image we took for the wheat kernels are presented in Figure S4. We use SmartGrain processed the same photo and compared the size estimation of SmartGrain with GridFree (Figure 5). We used the correlation between software estimated results and the manual measurements to examine the size measurement accuracies. Figure 5’ left figure shows the correlation of wheat kernel length between GridFree with manual measurement is R^2^=0.91, and the correlation between SmartGrain and the manual measurement is R^2^=0.90. The correlation between wheat kernel width estimation of GridFree with ground truth is R^2^=0.90, and the correlation between SmartGrain and the ground truth is R^2^=0.93. There is no significant difference in size estimation accuracies between the two software. The mean value of manually measured length is 6.35mm (*sd = 0.400*), and the width is 3.17mm (*sd=0.21*). The mean value of GridFree estimation for length is 6.21mm (*sd=0.422*), and 2.99mm (*sd=0.213*) for width. Mean value of SmartGrain estimation for length is 6.36mm (*sd=0.404*) for length, and 3.01mm (*sd=0.210*) for width.

**Figure 5.**
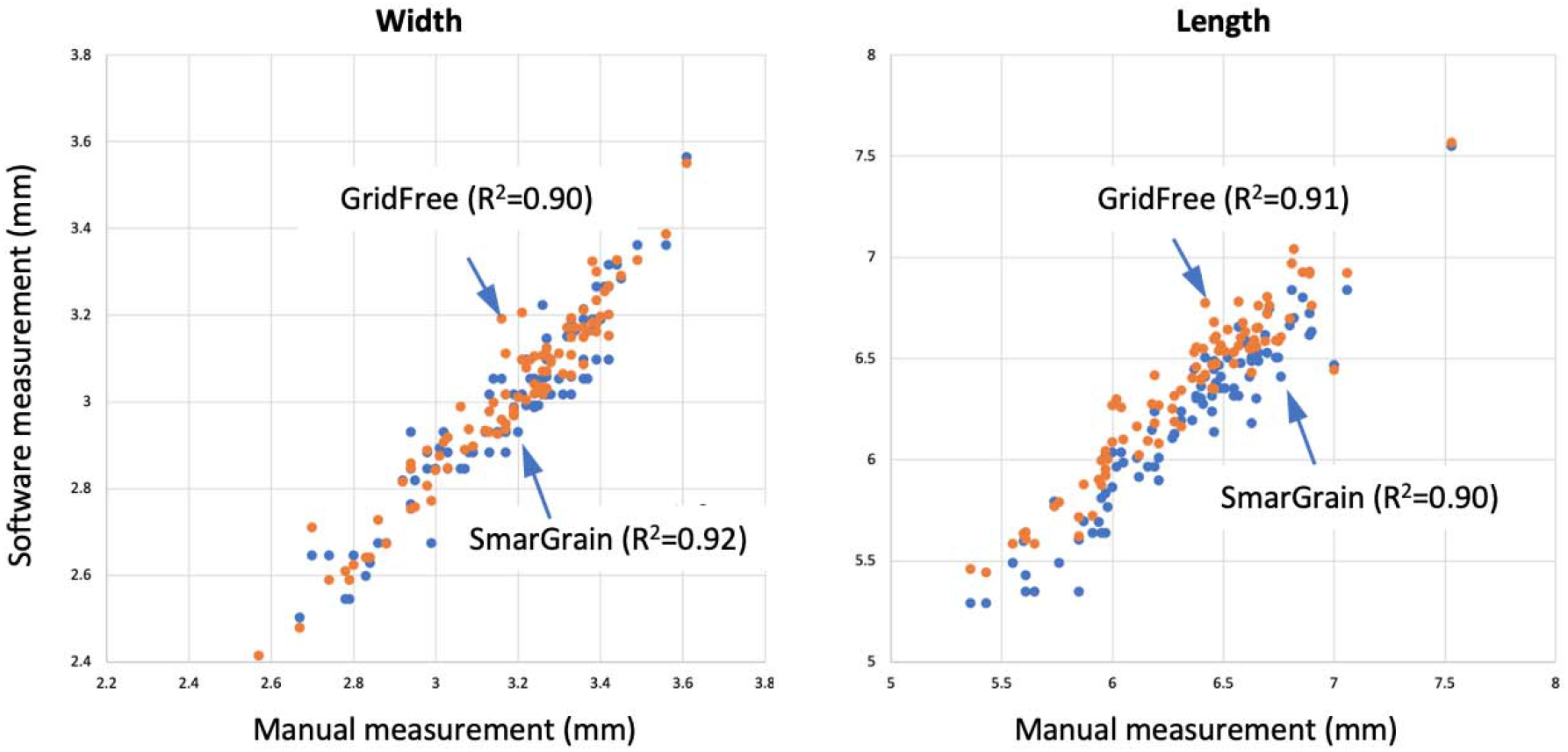
Comparison between manual measurements and predicts by GridFree and SmartGrain. A total of 92 wheat kernels were manually and compared with predicts by GridFree and SmartGrain on width (left panel) and length (right panel). GridFree has very similar accuracy as SmarGrain.

### Performances on counting and measuring variety type of objects

In addition to seeds (**Figure 6**), we found that GridFree can be used to count and measure spore balls from potato in an image provided by Tanaka Lab at Washington State University (https://labs.wsu.edu/tanaka-lab). The image was taken on a hemocytometer (**Figure 7**). The image (**Figure 7a**) was filtered at two levels to generate the image of POI, in this case, spore balls. The first level was the selection among the PCs derived from the three channels of the raw image and their 12 indices (**Figure 7b-d**). Of the 12 PCs, PC 6 best differentiated the spore balls from the other objects in the image, including the hemocytometer background and other particles (**Figure 7d**). The image based on the selected PC contains other patterns that are not visible to human eyes, such as the hemocytometer lines. After the second level of filtering by cluster analysis (**Figure 7e**), the hidden patterns in the selected PC images were separated (**Figure 7f** and **g**). One cluster represents the spore balls (**Figure 7h**). Segmentation on the image of POI was performed to ensure that the largest and the smallest segments matched the largest and the smallest spore balls. The final segmentation identified 12 spore balls in the image (**Figure 7i to l**).

**Figure 6.**
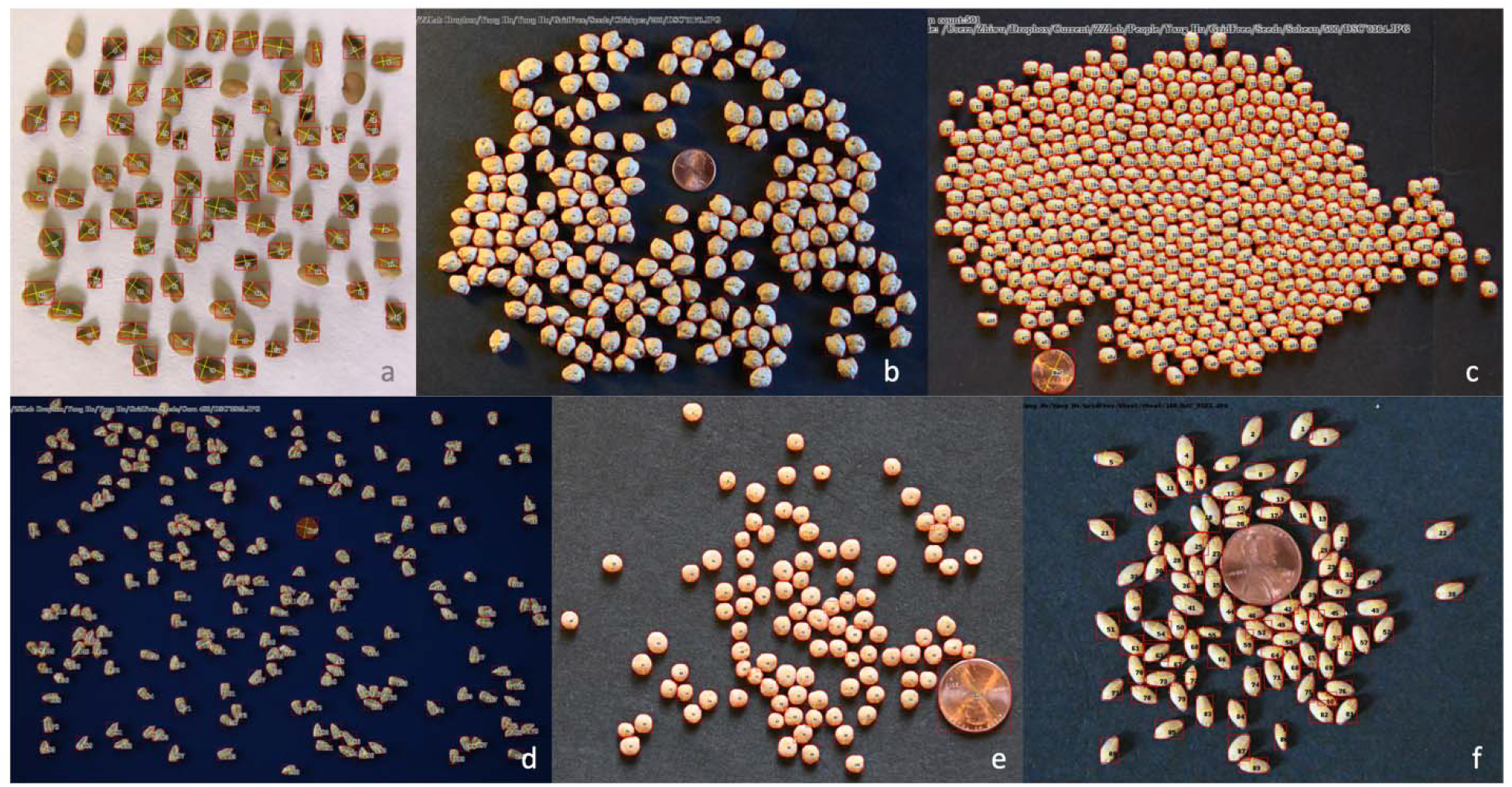
Segmentations of grain images by GridFree for multiple species. The segmentations are displayed as the frames of segmentation, the ordered numbers, and the size of segments in the aspect of length and width. The species include alfalfa (a), chickpea (b), soybean (c), corn (d), lentil (e), and wheat (f). The backgrounds colors were varied, such as white (a) and black (c). Each image contains a coin (US penny) as the reference.

**Figure 7.**
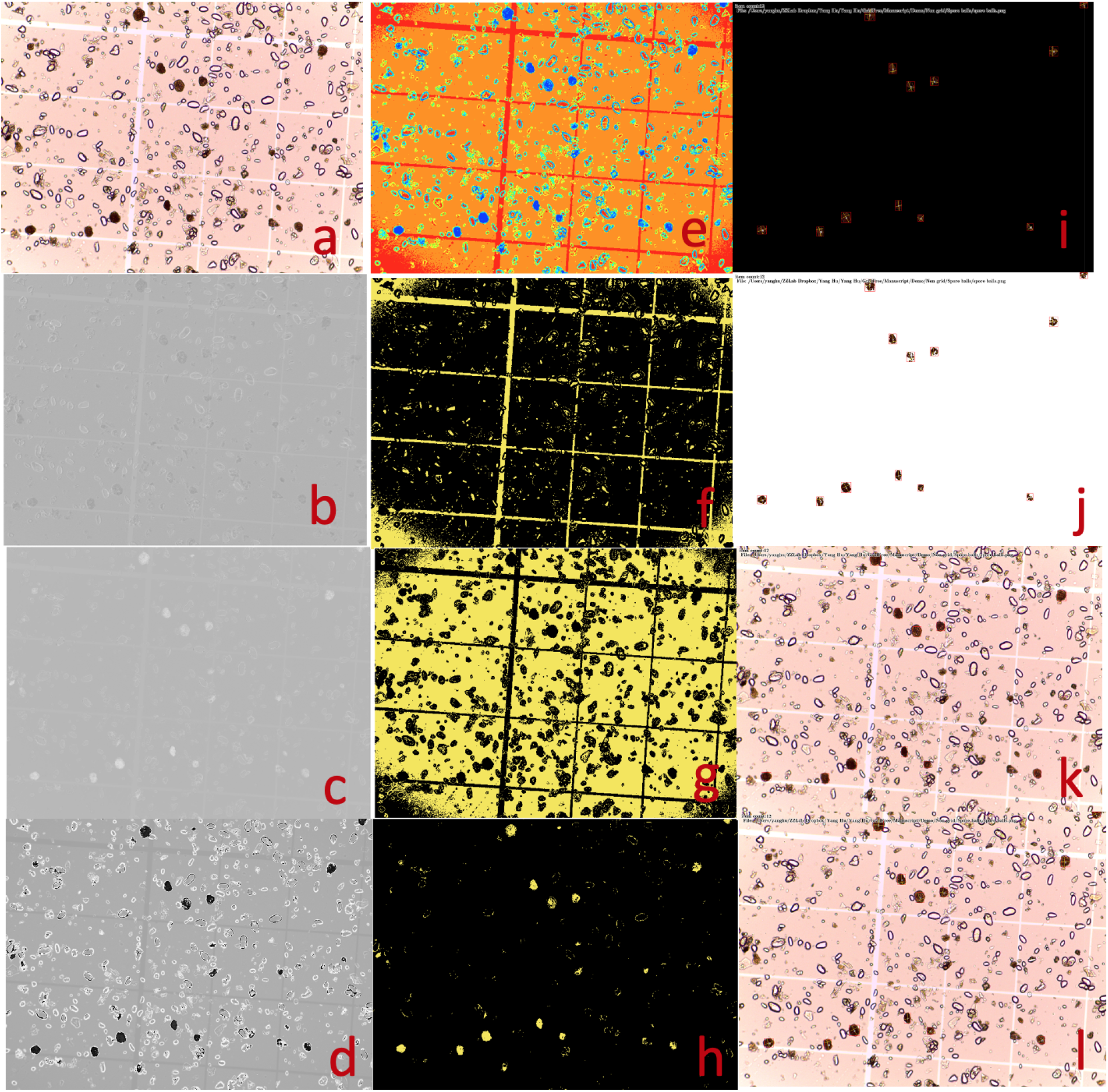
Counting and measuring spore balls from potato using GridFree. The picture was provided by Tanaka Lab at Washington State University (https://labs.wsu.edu/tanaka-lab). The objective was to count and measure the brown spore ball (a). Among the 10 Principal Components (PCs) from the RGB channels and the derived seven indices, three are displayed for PC1 (b), PC7 (c), and a combination of PC7 with RGB PC1 with weight of 0.312 (d). PC3 was the best at differentiating the spore balls from the background and was used for cluster analysis with five clusters (e). Among the five clusters, cluster 4 (f), 3 (g), and 1 (h) are displayed. Cluster 1 uniquely captured the spore balls and was used for segmentation. The segments were filtered to match the maximum and minimum spore ball sizes. The final segmentation results are displayed for extracted pixels of interest on a black background (i), on a white background (j), and on the raw image with segments represented as red-outlined boxes (l) and with measurements on length, width and counting numbers (k).

We applied the same process on a variety of images, ranging from cells and spores on a hemocytometer (**Figure S6**), E coli colony (**Figure S7**), chromosomes, antibodies, potato scabs, blood cells (**Figure S8**), to agricultural fields with different layouts, including orchards and plantation trees (**Figure S9**). Although the filtering of POI and segmentations were not fully optimized during the analyses for a large volume of images, the level of accuracy obtained suggests GridFree’s potential application in these different research areas.

## Discussion

In image analysis, successfully separating objects of interest from a noisy image background without explicitly examining a variety of color-indices is challenging. The discrepancy from normality of object size distributions adds even more complexity for counting and measuring seeds. Progress has been made through artificial intelligence, especially neural networks. For example, the software, Trainable Weka Segmentation (TWS) (Arganda-Carreras *et al.* 2017), can classify any pixel from a given image after enough training with a known dataset. The problem is that this training is also labor-intensive. GridFree was designed to fully capitalize on a human’s ability to make accurate judgments and to give the repetitive activities to computers. For the entire seed counting and measuring process, GridFree only requires users to decide which PCs and clusters most accurately represent objects of interest and to set expected area and size ranges of segments. Both of these judgments are facilitated through real-time previews of the segmentation results.

### Image decomposition using PCA

Color indices are commonly more effective than raw image channels such as RGB for separating objects of interest from the image background. Therefore, color indices are used by many software programs, including ImageJ, for users to select objects of interest (Rasband 1997). A variety of color indices are available, with some more similar than others (**Table S1**). However, making the decisions required to apply color indices is cumbersome and time-consuming. The difficulty is accentuated if users have little knowledge or experience using these color-index methods. To solve this problem, GridFree automatically conducts PCA on the raw image channels and 12 color indices (**Table S2** and **Figure S1-3**), and instantly displays outcome for users to make decisions.

For PCA, two methods can be used to derive eigenvalues and eigenvectors, which are then used to calculate PCs. One method uses covariance, and the other uses correlation among the variables. The variables are the raw image channels and color indices. The values of RGB channels range from 0 to 255, while the ranges for the 12 color-indices range from negative values to positive values, as the specific values vary in one image from another. We performed PCA on the three RGB channels and the 12 color-indices separately.

Users can select one from the two RGB PCs with the highest variance. The number of PCs is as same as the number of variables of color indices. The first PC contains more information than the second, the second PC more than the third, and the last PC contains the least information. Therefore, in GridFree, the first component of the color index is selected by default. Unlike the original variables of RGB channels that are most likely correlated, the 12 color indices PCs are independent of each other. In other words, after PCA, the original image is decomposed into independent PC images. Users can change the default if they find other PCs better represent the objects of interest. With the consideration of using information from RGB channels, users can make a combination of the RGB PCs with the 12 color indices by changing RGB PCs’ weights.

### Selecting the levels of PCs using K-Means clustering

For the selected PCs, GridFree applies an unsupervised machine learning method, K-Means Clustering, to classify different ranges of the values of PCs into groups. K-Means can be initiated by dragging the clustering bar away from 1 to a specific number of clusters. Each clustering group is represented by a color and displayed on the display panel to assist users in identifying the objects of interest. Users can check the corresponding checkboxes to define the objects of interest. A binary image is created with pixels in the selected clusters coded as 1 for the objects of interest, and the rest of the pixels as 0 for the background.

### Interactive dynamic segmentation

The existing seed counting applications (GrainScan and ImageJ) (Rasband 1997; Whan *et al.* 2014) rely on an initial inquiry to users for kernel size ranges, which is a difficult question to answer. GridFree solves this problem by cross-referencing displays of the segmentation image and the graphic distribution of segment sizes. That is, the preliminary segmentation plot and segmentation image are linked, allowing users to click on any data point in the plot and simultaneously view its corresponding segment on the display panel (flashing yellow rectangle). When grain kernels are closely adjacent, they can be mistaken as one item with dimensions larger than single seeds and then depicted as outliers. In contrast, over-segmentation will result in multiple data points assigned to parts of single seeds and/or noise artifacts assigned as kernel pixels, thus forming another type of outlier. To correct these errors, GridFree employs interactive, dynamic thresholding with four click-and-drag bars that help users set area and size thresholds to further refine the segmentation. Multiple iterations may be necessary before reaching the most accurate segmentation result.

### Candidates for reference objects

GridFree tolerates multiple shapes and sizes of the reference object; however, the reference should be included as part of the selected clusters of interest so that their colors match. Otherwise, the reference will not be visible on the binary image. When the binary image cannot wholly account for the complete reference, the grain kernel area and size estimates will be smaller than the actual dimensions. Users should be cautioned to select the PCs that ensure the entire reference surface is recognized. One tip is to choose reference objects with colors similar to objects of interest, giving the reference a greater chance of being included during the selection of objects of interest.

### Seed measurements

GridFree provides measurements of length and width in addition to the total area. The differences between length and width vary from species to species. Some species have very different scales for length and width, such as wheat, corn, and alfalfa. Round objects, such as soybean and lentil seeds, exhibit minimal differences between length and width. The stage of maturity also affects shape. For example, fully mature alfalfa seeds tend to be longer than less mature seeds. For seeds with two pointy ends, such as wheat, length and width measurements by GridFree closely match the morphological measurements. However, when seeds such as corn have one flat end, a substantial proportion of that length is not measured at the center of kernels.

We did not make comparisons with other software programs on length and width for two reasons. First, only a limited number of software programs can measure these characteristics. Second, obtaining good standards to evaluate these measurements is not trivial. Measuring these characteristics manually is time-consuming and subject to human error and inconsistencies; therefore, the measurements may not be accurate enough to serve as standards. The confirmation of SmartGrain was conducted by the association study between the digital measurements and the genetic markers covering the whole rice genome. The identified genetic loci served as evidence that the digital measurements were useful (Tanabata *et al.* 2012). Instead of focusing on one species for measuring seed length and width, our study examined a variety of species, including wheat, soybean, lentil, chickpea, black beans, canola, alfalfa, and corn.

For seed counting, we made the comparisons between ground truth measurements and the counts using GridFree and the other four software programs. The ground truth was designed to have 100, 200, 500, and 1000 seeds for four types of grains (black bean, canola, chickpea, and lentil. For the alfalfa experiment, the design had four levels: 100, 200, 400, and 1000 kernels. The advantage of counting by images is that the counting results can be repeatedly examined for errors. Especially, human errors for manual counting are difficult to identify unless counting by images is used for verification.

### Limitations

Multispectral images have gained popularity recently. GridFree only processes the first three channels as RGB and derives 12 color indices accordingly. Incorporating more channels for the raw images could be done without much difficulty. Currently, users can combine one from the 12 color indices PC with one from the first two RGB PC with weight ranges 0.0 through 1.0, with an interval of 0.002. The price that users pay is having to drag the sliding bar of RGB PCs and selecting color indices PC images.

Size reference object function is useful in single image mode but is disabled under the Batch mode. To make the results comparable among images in the same batch, these images must be taken with the same distance from cameras to the objects. To make images comparable between batches, users need to process one image, then use the process parameters to process the rest images. If image contains reference objects, then users have to process all the images manually that to specify reference for images in the batch. The measurements based on pixels must be converted accordingly.

Although GridFree can be used for segmentation of images with grid patterns, such as field plots for agricultural experiments (**Figure S9**), difficulty arises when assigning the identification for the segments. A mistake on a single segment will cause mismatched identifications for the subsequent segments, requiring manual correction. GridFree does not include a method to set particular numbers of rows and columns as other software packages do. For example, for images with grid patterns, GRID (Chen and Zhang 2020) can manually set the number of rows and columns, which limits mismatching across rows or columns in a situation of mistaken segmentation. In fact, the GridFree package was motivated by GRID to analyze images without the requirement of grid patterns, hence the name, GridFree.

GridFree does not have functions to flatten skewed image, one reason is flattened image usually changes shapes and length-width ratios of items. Another reason is adjusting a skewed image requires the image contains boundaries, which limit the image environments.

## Conclusion

We developed a two-level filtering method for pixels of interest in photographic image analysis, plus a method of filtering segments using both segment size and number of pixels of interest. We implemented the two methods in an open-source Python package, GridFree, with a friendly GUI. The software can count grain kernels in photographic images with higher accuracy than existing software packages. With GridFree, users are completely liberated from manually counting seeds at high accuracy. All the processes are automated and only require users to make judgments to ensure the outcomes accurately segment the original input image, based on previewing preliminary segmentation images. GridFree also provides measurements for the segmented objects, including area, length, and width. In addition to seeds, the software has the potential for counting and measuring other types of objects, such as cells, E Coli colonies, spores, agricultural field plots, and orchard or plantation trees.

## Materials and Methods

We developed methods and a software package to select pixels corresponding to objects of interest without explicitly examining a variety of color-indices. Furthermore, our methods allow users to conduct segmentation without the restriction of normality of segmentation size to separate objects adjacent to each other. PCA is employed to decompose the raw channels of the input image and their derived color-indices and display as component images. The input image is classified pixel-wise based on the selected PCs using K-Means cluster analysis. The selected clusters are used to define POI and group all other parts of the image as background to form a binary image. To overcome the restriction of normality of item pixel sizes, we developed a dynamic divide-merge procedure combined with automatic segmentation using Breadth-First Search (BFS) and watershed algorithms jointly. The following subsections include further details about image pre-processing, image segmentation algorithms, dynamic threshold setting, and the divide-combine strategy for final segmentation.

### Image decomposition using principal component analysis

Color index approaches are applied to the raw RGB channels, and 12 color-indices are derived, including PAT_R, PAT_G, PAT_B, DIF_R, DIF_G, DIF_B, ROO_R, ROO_G, ROO_B, GLD_R, GLD_G, and GLD_B (**Table S1**). We examined 30 color indices derived from raw RGB channels via a heatmap (**Figure S1**) and noticed the 12 indices have the least correlation with each other across channels. RGB channels and the 12 color-indices are represented in matrices (A), with rows equal to the product of h and w and number of columns equal to the sum of c and d. The h and w represent the height and width of the image; c and d represent the number of channels and color-indices. GridFree applies PCA to the first three columns (RGB) and the last 12 columns (indices) in Matrix A separately to derive the RGB PCs and index PCs, and stores them as matrix B. Correlation is used instead of covariance to ensure that the contributions of columns (raw channels and color-indices) do not depend on their scales.

### K-Means clustering on selected principal components

By default, the first color index PC is selected. The selected index PC can be averaged with either RGB PC1 or PC2, displayed with grayscale, and are used to conduct pixel-wise cluster analysis using an unsupervised machine learning method, K-Means clustering. Pixels are displayed in colors corresponding to their clusters. The selected cluster (by users) is coded as value 1 and the rest as 0 to form a binary image with a height of h and width of w.

### Dynamic segmentation

We used BFS and watershed algorithms jointly as the essential building blocks for segmentation. BFS is an algorithm to explore nodes and edges of a graph, particularly useful for finding the shortest path. Applications include finding connected components of a graph. Two nodes are adjacent pixels if an edge is between them. A segment is a set of nodes connected with edges. We used BFS to initiate segmentation. The watershed algorithm is also commonly used for segmentation based on the representation of grayscale images as topographic relief with watersheds dividing different basins. Euclidian distance was used to choose markers, which are local minima of the image of the selected PCs. The basins are flooded from the bottom, the marker, to the top until basins attributed to different markers meet on watershed lines.

BFS has an advantage over the watershed algorithm for an image filled with grain kernels. In such a case, the watershed algorithm tends to partition single objects without smooth surfaces such as wrinkled kernels. That is, the watershed algorithm will label each side of a wrinkled kernel as a separate kernel. Conversely, the watershed algorithm has an advantage over BFS for images containing adjacent kernels. We used the watershed algorithm to deal with the image portion containing a group of adjacent kernels. The limitation of the watershed algorithm is the difficulty in locating center points for those adjacent kernels whose rims are smooth, with no visible dent.

To overcome the limitation of the watershed algorithm, we applied a dynamic, search-region method because the first step of the watershed algorithm is to identify markers from an image. The number of markers decides the number of objects after segmentation. We initialize the search region as the mean area, in pixels, of correctly segmented objects. The possible number of objects after segmentation can be estimated by dividing the total area of adjacent objects by the average area of segmented objects. If the number of markers is less than the estimated value, the search region shrinks by one-half of the mean area. The process iterates until the number of markers equals the estimated value or the search region reduces to one.

To separate adjacent objects whose rims have dents, we applied a boundary shrinkage method to identify the center of each object. For segments above the upper boundaries defined by users, boundary shrinkage is applied to reduce the segments. Boundaries are the pixels with at least one background pixel neighbor. Boundary pixels are iteratively disabled as background pixels until the segment satisfies the upper bounds, or divides into two or more segments. The new segments are recursively processed to satisfy the upper bounds. For any pixel p in the area, the method searches pixel p’s eight neighbor pixels. If any of the neighbors belong to the background, then pixel p will be labeled as the boundary. The neighbor pixels’ coordinates are obtained from adjusting pixel p’s coordinate values by +/− 1 to the current pixel location’s x and y coordinate values eight times, indicating pixel p’s eight neighbors. All pixels labeled as boundary will be temporarily disabled from the area of interest. The removed pixels will be re-enabled once the boundary shrinkage is completed.

A segment resulting from the initial segmentation is permanently removed if its area in pixels or size as diagonal length is below the lower bounds defined by users. The completion of object segmentation may result in new segments below the lower bounds. These new segments will go through a boundary expansion. Such a segment will merge with its neighbor segment. Specifically, the merging process will create a new segment that satisfies the lower bounds. The expansion stops when no disabled neighbor can be found. A segment is deleted if the expansion still does not satisfy the lower bounds.

### Implementation in GridFree software package

We designed the GridFree user-interface to have three panels, with the main panel of display on the left, the filter panel on the right, and the control panel on the bottom. There are five mode for the main panel of display, corresponding to the raw image, PC image, the cluster image, the binary image of selected clusters, and the output image of segmentation. The filter panel contains two sliding bars, an array of checkboxes, and a filter display. The first sliding bar controls the weight of RGB PC1 or PC2 over the selected index PC. The contents on the filter display change according to mode of the main panel of display. Under the mode of selected PCs, the filter display demonstrates the 12 thumbnails of the index PCs. Clicking the thumbnail magnify the corresponding PC on the main panel of display. For a selected index PC, moving the RGB PC sliding bar away from the center will increase the weight of PC1 toward the left, or PC2 toward the right. The main panel of display panel will update soon after users move the cursor out of filter panel.

The second sliding bar controls the number of clusters for K-Means clustering analysis. The checkbox array accompanies each cluster for selection. The selected clusters are displayed as a binary image with the rest of the clusters as the background. Below the sliding bars and checkboxes is the filter display. The cluster image and the binary image are displayed by the main panel of display and the filter display respectively. Their displaying places can be switched by choosing the display mode (Cluster and Selected) on the main panel.

When the main display panel is at output mode, the filter panel displays segment distribution of area against size using the Canvas class of the Tkinter Python library. The horizontal axis represents the area as number of pixels in each segment. The vertical axis represents the size of the segments as their diagonals. The scatter plot has four moveable bars for dynamic threshold setting. The two vertical red bars are used to set the upper and lower thresholds for the object area. The two horizontal blue bars are used to set the thresholds for object size. The four bars are controlled with a mouse or similar computer pointing device using a click-and-drag process. Users move the bars by clicking and dragging them to the desired positions. The thresholds become guidance for segmenting adjacent grain kernels, using a divide-and-combine strategy.

### Cross-reference display of segmentation

For the purpose of facilitating the positioning of the threshold bars, user-selected segments are cross-referenced on both the image in the main display panel and the scatter plot display in the filter panel. The selected segments are highlighted (flashing yellow). When a segment is selected on the main display, its corresponding dot in the scatter plot of the filter panel is also highlighted, which helps users determine the proper thresholds. Conversely, when a dot is selected on the filter display, the highlighted cross-referenced segment on the main display panel help users determine the validity of the segments, especially for the outliers.

### Candidates of reference objects

To convert grain kernel area and size from pixels to physical units such as millimeters, GridFree uses the ratio of the reference object area in number of pixels to the actual area of the reference in millimeters. This value is then used to convert area and size in pixels to area and size in millimeters. This method tolerates types and shapes of size references, and users can choose any object with a known area as a size reference. Only the total area matters.

### Batch mode

Pictures taken under similar environments usually share similar features. Such properties facilitate GridFree batch processing by using a set of parameters. These parameters include PC selection, RGB PC weights, k value for K-Means, cluster selection, and the upper and lower bounds for segment area and size. Batch mode is activated by checking the “Batchmode” checkbox. To specify the location of the input images, where to export results, what parameters to use, and when to start, we added additional functions for the command buttons corresponding to single image mode. The Image and Export buttons tell the user where the input images are located and where to export results, respectively. The Pilot Button imports the parameter file. The parameter file has the same format as the output of parameter file for the single image mode. The details are included in the user manual. To start the batch mode process, click the Segment button. Batch mode processes images one after another. The reference object is disabled under batch mode.

**Table 1.** Characteristics of software packages for counting and measuring grain kernels.

| Name | Free to public | Platform | First release | Latest update | Reference |
| --- | --- | --- | --- | --- | --- |
| SmartGrain | Yes | Windows OS | 2012 | 2012 | (Tanabata et al. 2012) |
| GrainScan | Yes | Windows OS | 2014 | 2014 | (Whan et al. 2014) |
| SeedCounter | Yes | Android | 2017 | 2017 | (Komyshev, Genaev, and Afonnikov 2017) |
| MUSeed | NA | Android | 2017 | 2017 | (Gao et al. 2017) |
| GainTKW | NA | Android | 2018 | 2018 | (Wu et al. 2018) |
| ImageJ | Yes | iOS/Windows | 1997 | 2019 | (Rasband and others 1997) |

## Acknowledgement

This project was partially supported by the USDA National Institute of Food and Agriculture (Hatch project 1014919, Award #s 2016-68004-24770, 2018-70005-28792, and 2019-67013-29171), the National Science Foundation (Award number DBI 1661348), and the Washington Grain Commission (Endowment and Award #s 126593 and 134574). The authors thank Meijing Liang and Wanling Li for testing and advising the software, and Linda R. Klein for helpful comments and editing the manuscript.

## Notes

### Competing Interest Statement

The authors have declared no competing interest.

