## Supplement figures for "GridFree: A Python Package of Image Analysis for Interactive Grain Counting and Measuring"

**Supplementary**

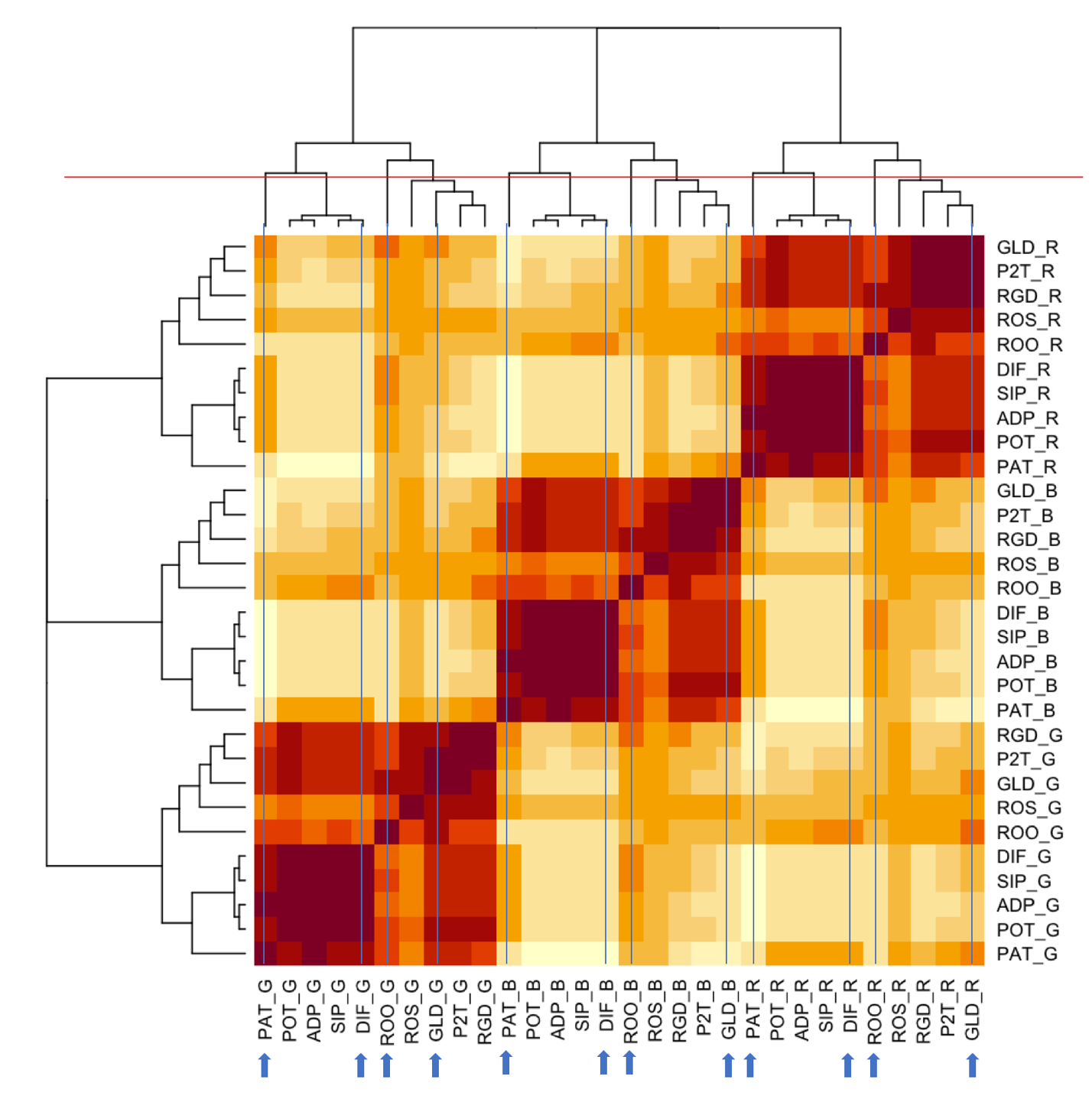

**Figure S1. Relationship among the 30 color indices derived from three channels.** The three channels (RGB) were independently simulated from a uniform distribution between 0 and 255. The 30 indices are clearly classified into three groups according to the three channels. For each channel, the cladograms were cut at the level to form four branches (red line). There are six branches with a single-color index that belong to the three categories consist of PAT, DIF, or ROO. For each of the remaining branches with multiple color indices, the color index was selected if it belongs to the three categories or GLD. In total, 12 color indices were selected corresponding to three channels and four categories. These indices have the least correlation with each other across channels.

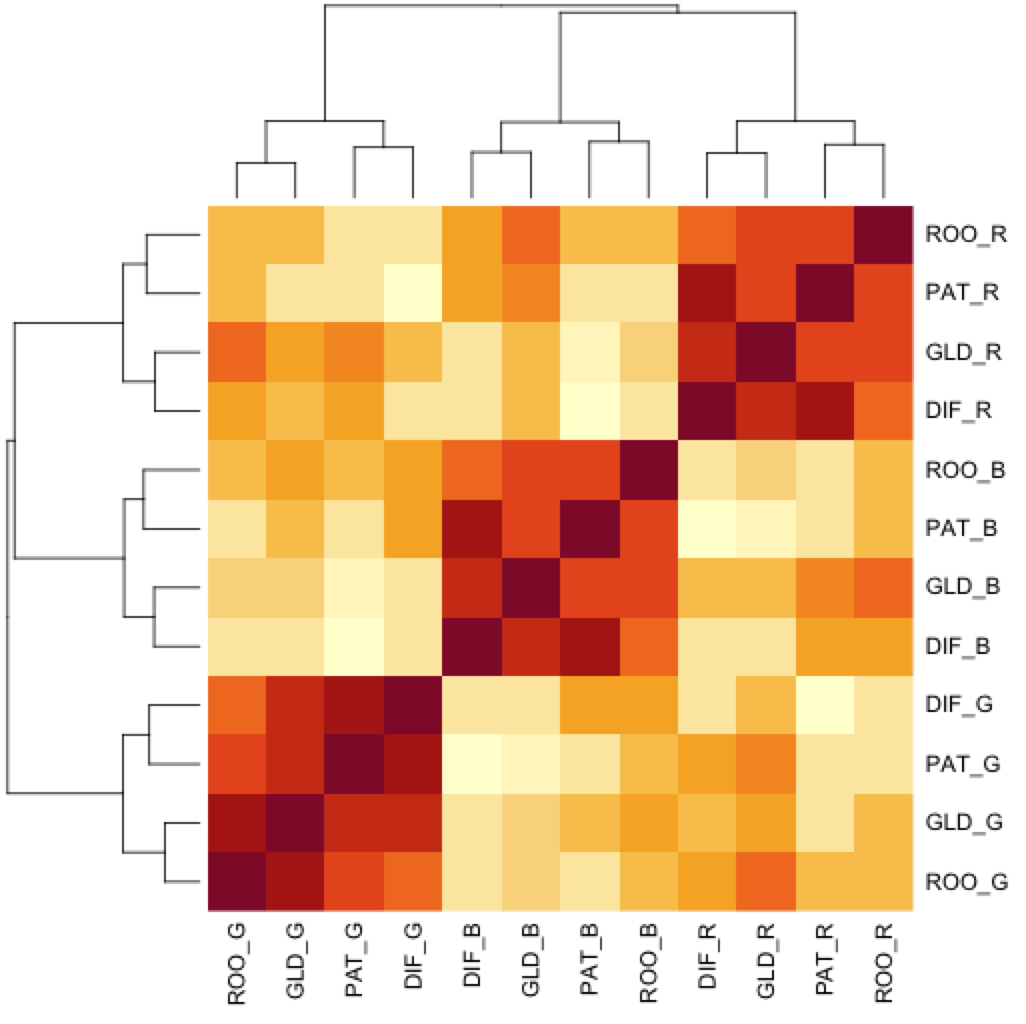

**Figure S2. Relationship among the 12 color indices derived from three channels.** The three channels (RGB) were independently simulated from a uniform distribution between 0 and 255. The 12 indices are clearly classified into three groups according to the three channels. The correlations among the indices within colors are much higher than the ones among the colors. The dark red indicates the Pearson correlation coefficient of 1 and white as zero.

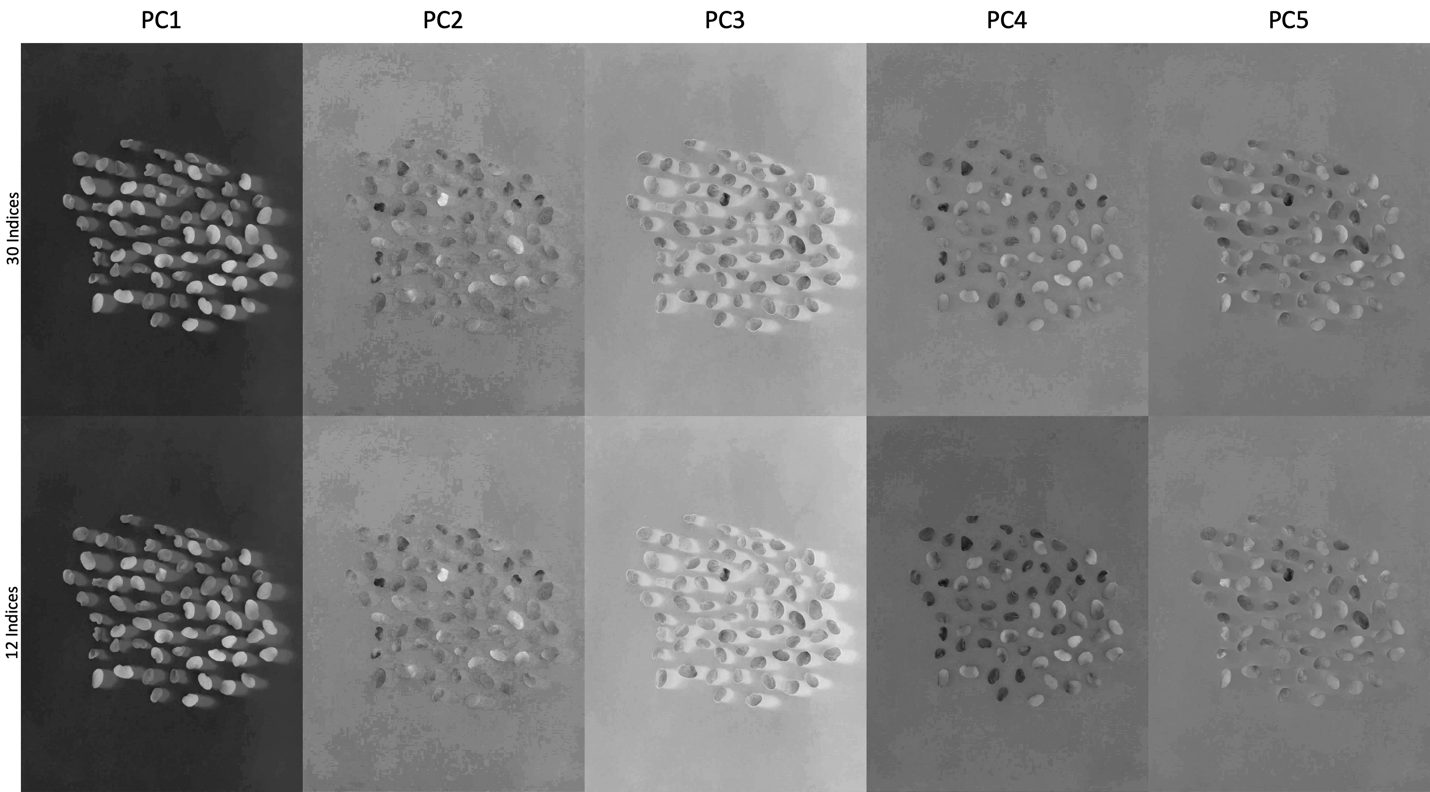

**Figure S3. Information preservation of the 12 color indices selected from 30 color indices.** The color indices were calculated on an image of alfalfa seeds using three channels (RGB). Principal component analyses were conducted on the 30 color indices and the 12 selected separately. The first five Principal Components (PCs) are displayed in gray scale at the top panel for the PCs derived from 30 indices, and bottom panel for the PCs derived from the 12 selected indices. The 12 selected indices preserve almost all the information from the 30 indices.

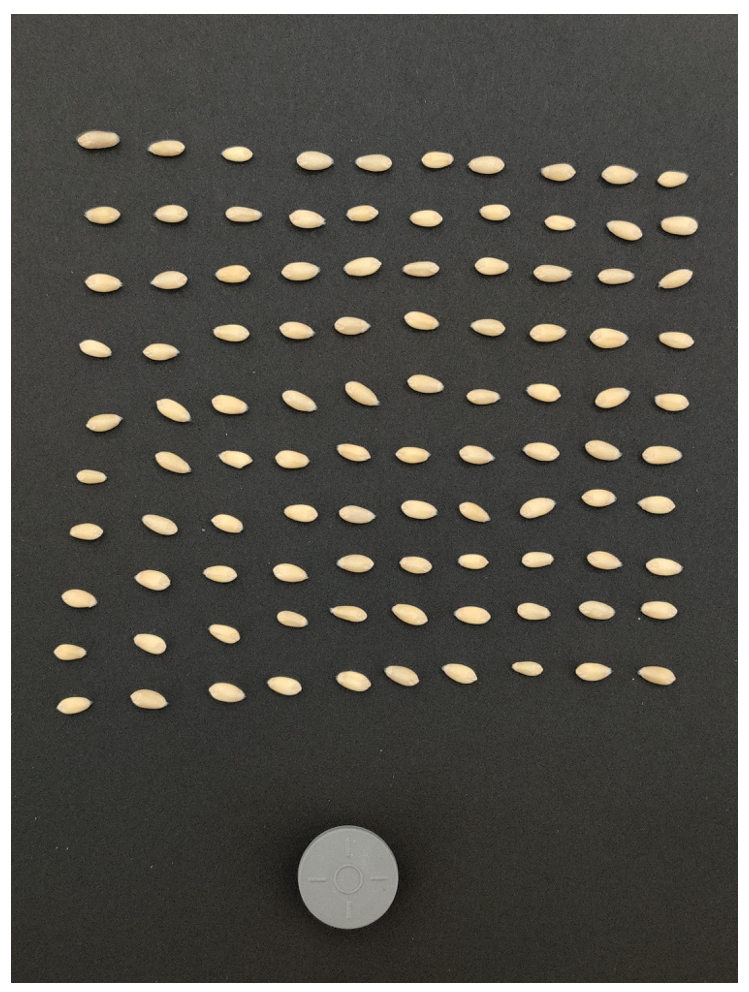

**Figure S4. Image for wheat kernel size validation**

The image was taken by a OnePlue Pro7 Android phone, with regular photographing setting. The image resolution is 3000x4000. A spherical item at the bottom is considered as size reference for size estimation for GridFree and SmartGrain.

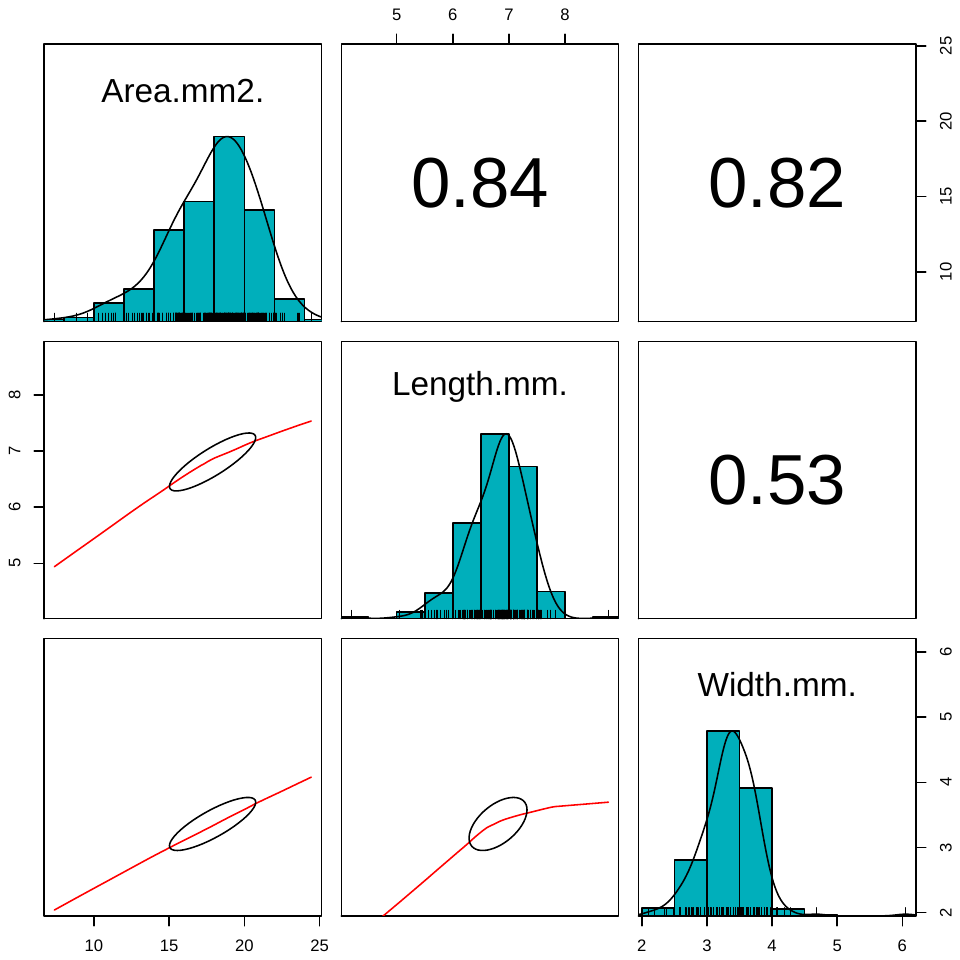

**Figure S5. Distributions, scatter plots, and correlations among area, length, and width for wheat kernels.** The distribution of areas, lengths, and widths of wheat seeds are displayed on diagonal (units in millimeter). The correlations between area, length, and width are displayed as scatter plots in the lower triangle; correlation coefficients are displayed in the upper triangle.

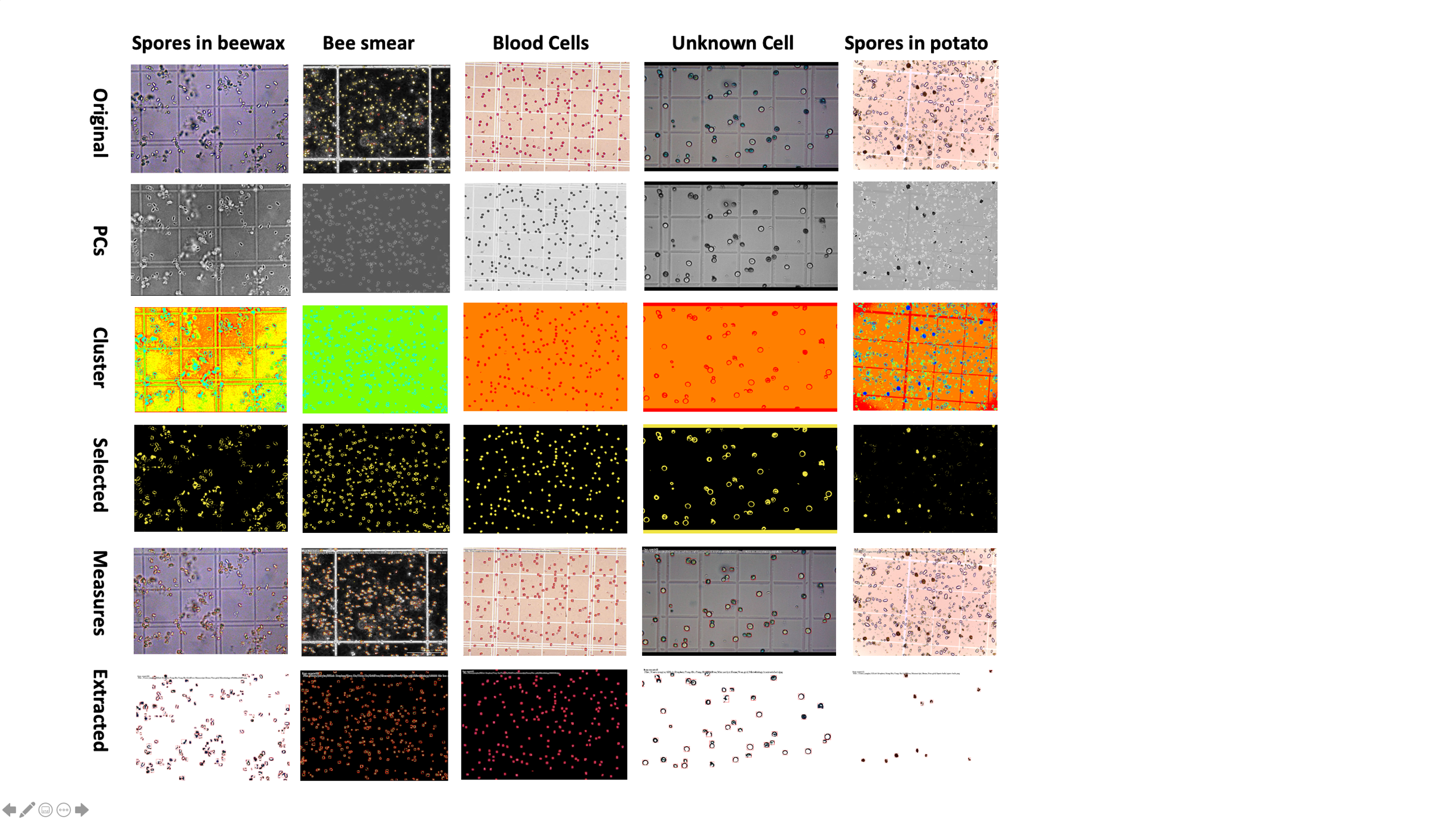

**Figure S6. Spores and cells on hemocytometer.** Five types of objects on hemocytometers (row-wise) were counted and measured. The original images, measurements, and intermediate processing images are displayed column-wise.

Image source:

Spores in beewax: <http://pds61.cafe.daum.net/image/6/cafe/2008/02/15/22/30/47b593ea15d07>

Bee smear: <https://4.bp.blogspot.com/-6eyv7GbWd78/VUKBZVXDZ-I/AAAAAAAAGbs/kQY5GZl0ym0/s1600/150326%2B20x%2Bbee%2Bsmear-2-analysis.tif>

Blood cells: <https://www.wardsci.com/stibo/low_res/std.lang.all/61/53/25306153.jpg>

Unknown cells: <https://i.ytimg.com/vi/50SwIYkDxEQ/maxresdefault.jpg>

Spores in potato: Provided by Tanaka Lab at Washington State University (<https://labs.wsu.edu/tanaka-lab>)

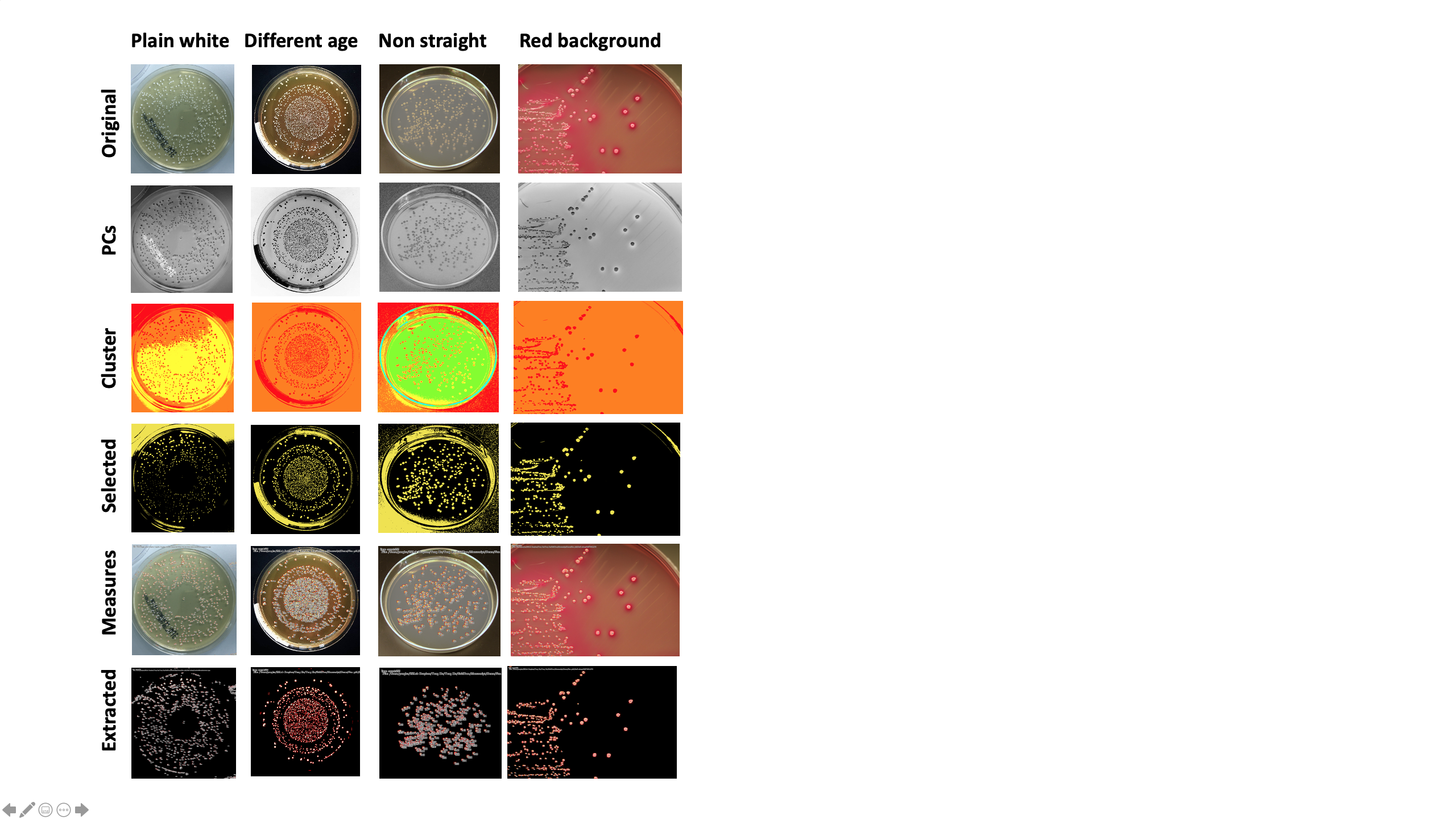

**Figure S7. E coli colony with different background and angles.** E coli on four types of backgrounds (column-wise) were counted and measured. The original images, measurements, and intermediate processing images are displayed row-wise.

Image source:

Plain white: <https://mltgeeks.com/wp-content/uploads/2018/09/Lactobacillus-plantarum-1.jpg>

Different age: <https://1.bp.blogspot.com/-JJxBcri0bgc/WRwWyJ0qcSI/AAAAAAAAAUc/PcFhUSm3BwUL14dRS53TO-9hokPZq0E9ACLcB/s320/1799851_518039825010535_2086813812_n.jpg>

Away from the top (non straight): <https://i.pinimg.com/originals/1f/d5/fa/1fd5fa1765e5681634bb9ba260780f50.png>

Red background: <https://4.bp.blogspot.com/-peNjHGbf7XQ/WdpwnoEbzzI/AAAAAAAAAbY/qHLmG563ejcaix_iayjYO3EgmSGig8xBQCLcBGAs/s1600/IMG_2276.JPG>

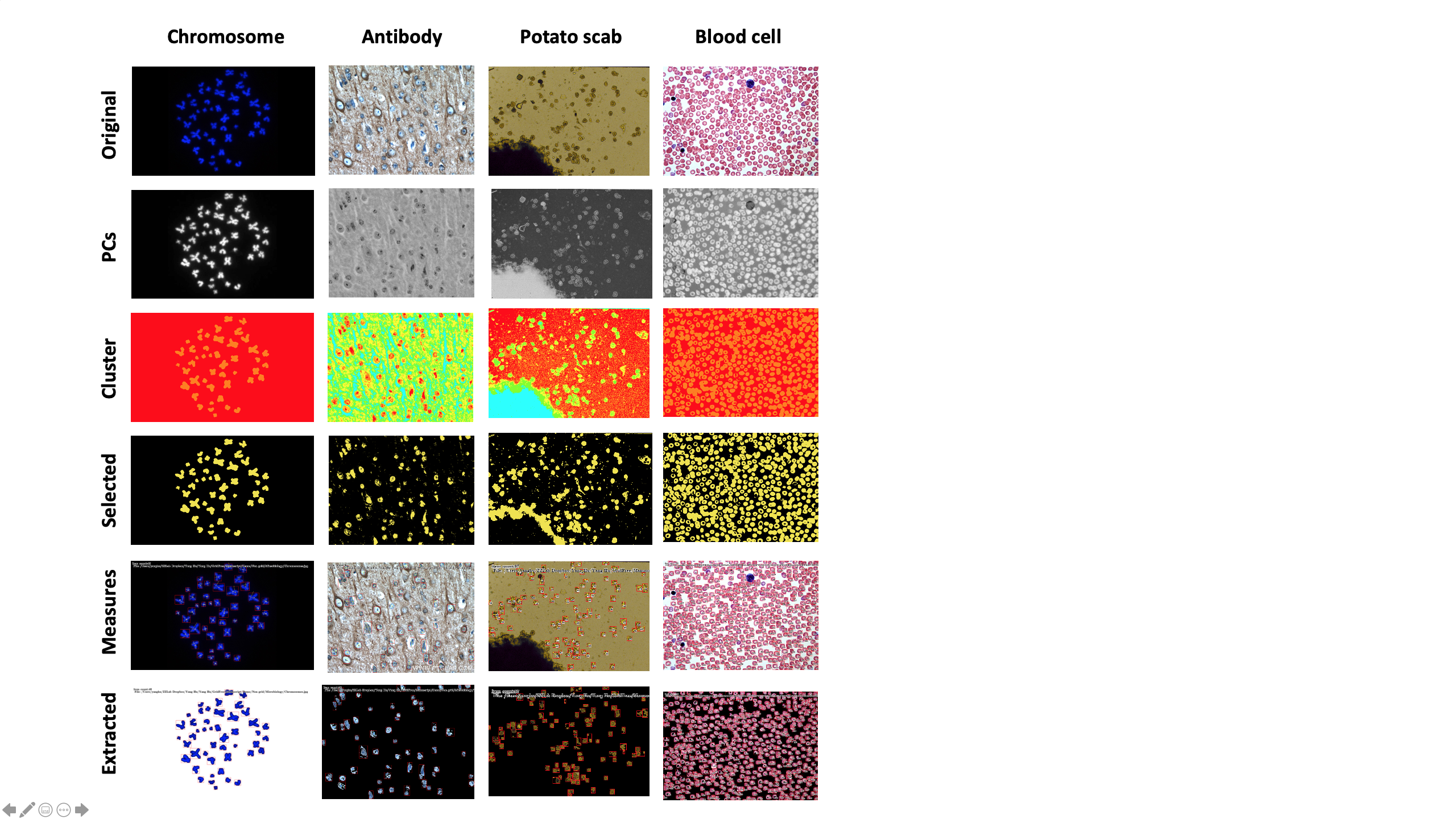

**Figure S8. Counting under microscope.** Chromosomes, antibodies, scabs, and blood cells (column-wise) were counted and measured. The original images, measurements, and intermediate processing images are displayed row-wise.

Image source:

Chromosome: <https://membs.org/membs/uploads/news_images/m.jpg>

Antibody: <http://www.ptglab.com/Products/Pictures/TUBB3-Antibody-10068-1-AP-IHC-18257.jpg>

Potato scab: <https://pnwhandbooks.org/sites/pnwhandbooks/files/plant/images/potato-solanum-tuberosum-powdery-scab/cystoriapowderyscab.jpg>

Blood cells: <https://paramedicsworld.com/wp-content/uploads/2017/12/PERIPHERAL-BLOOD-SMEAR-DIFFERENTIAL-LEUCOCYTE-COUNT-DLC-PBS-PERIPHERAL-BLOOD-FILM-THIN-BLOOD-SMEAR-PERIPHERAL-BLOOD-FILM-BLOOD-SMEAR-BLOOD-FILM.jpg>

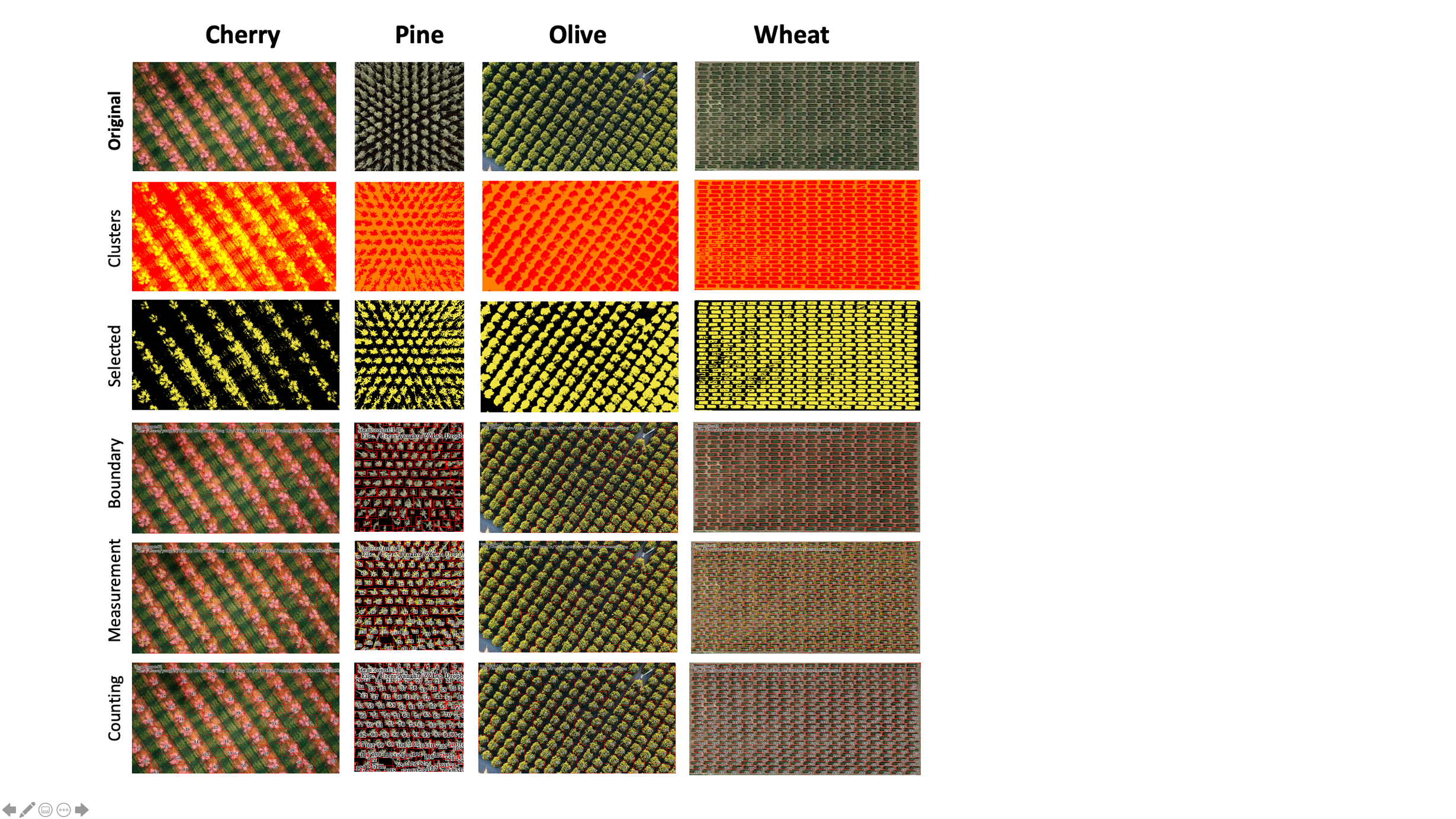

**Figure S9. Counting and measuring field plots.** Cherry, pine, olive, and wheat plots (column-wise) were counted and measured. The original images, measurements, and intermediate processing images are displayed row-wise.

Image source:

Cherry: <https://i.pinimg.com/474x/62/bf/83/62bf836d86e527d881379c62c1611dff--peach-trees-peach-orchard.jpg>

Palm: <http://cdn.shutterstock.com/shutterstock/videos/11359739/thumb/1.jpg?i10c=img.resize(height:160)>

Pine: <https://media.wired.com/photos/5a55d9884bcfdd7311969492/1:1/w_1000,h_1000,c_limit/Baumschule-016.jpg>

Olive: <https://www.oliveoiltimes.com/business/drones-olive-farms/58796>

Wheat: Google satellite image on Spillman Agronomy Farm at Washington State University (46.695693, 117.149742)

**Table S1. Details of color-index approaches***

| Name | Description | Equation | Applications |
| --- | --- | --- | --- |
| NDI | Normalized difference index | 128*(G-R)/(G+R+1) | (Meyer and Neto 2008) |
| Greenness (GCC) | Green chromatic coordinate | G/(G+R+B) | (Woebbecke et al. 1995) |
| VEG | Vegetativen | G/(R^(0.667)*B^(1-0.667)) | (Wan et al. 2018) |
| CIVE | Color Index of Vegetation Extraction | 0.44*R+0.811*G+0.385*B+18.7845 | (Kataoka et al. 2003) |
| MExG | Modified Excess Green | 1.262*G-0.844*R-0.311*B | (Hamuda, Glavin, and Jones 2016) |
| NDRB | Normalized difference of primary and pigments | (R-B)/(R+B) | (Kawashima and Nakatani 1998) |
| NGRDI(GDVI) | normalized difference green/red index | (G-R)/(G+R) | (Tucker 1978) |

* Red, Green, and Blue are three channels in RGB images.

**Table S2. Definition of 30 color indices derived from RGB channels***

| Index | Description | R primary | G primary | B primary | Applications |
| --- | --- | --- | --- | --- | --- |
| ADP | Average Difference Proportion | (2*R-G-B)/  (2*R+G+B) | (2*G-B-R)/  (2*G+B+R) | (2*B-R-G)/  (2*B+R+G) | (Tucker 1978) and (Meyer and Neto 2008) |
| DIF | DIFference | 2*R-G-B | 2*G-B-R | 2*B-R-G |  |
| GLD | GoLDen ratio | R/(B^0.618*  G^0.382) | G/(B^0.618*  R^0.382) | B/(G^0.618*  R^0.382) | (Wan et al. 2018) |
| P2T | Proportion to Total of others | 2*R/(G+B) | 2*G/(R+B) | 2*B/(G+R) |  |
| PAT | Proportion Among Two bands | R/(R+G) | G/(G+B) | B/(B+R) |  |
| POT | Proportion Of Total | R/(R+G+B) | G/(R+G+B) | B/(R+G+B) | (Woebbecke et al. 1995) |
| RGD | Reverse Golden Ratio | R/(G^0.618*  B^0.382) | G/(R^0.618*  B^0.382) | B/(R^0.618*  G^0.382) | (Wan et al. 2018) |
| ROO | Ratio Over Other | R/G | G/B | B/R |  |
| ROS | Ratio Of Square | R*R/(G*B) | G*G/(B*R) | B*B/(R*G) |  |
| SIP | Square Increase Proportion | (R*R-G*B)/  (R*R+G*B) | (G*G-B*R)/  (G*G+B*R) | (B*B-R*G)/  (B*B+R*G) |  |

* Red, Green, and Blue channels are indicated as R, G, and B, respectively.

**Appendix**

**Image pre-processing method:**

1. Centralize matrix A, Centr(A)
2. Obtain Correlation coefficient matrix of transposed centralized matrix A, Corr(Centr(A^T^))
3. Obtain eigenvector and eigenvalues from Corr(Centr(A^T^)), Eig(Corr(Centr(A^T^)))
4. Rank eigenvectors by eigenvalues
5. Obtain Cauchy product of centralized matrix A, Centr(A) and Eig(Corr(Centr(A^T^))), which is the matrix B

**The pseudo-code of BSF application for GridFree:**

Input: a graph of target component pixels

Output: a labeled graph of target component pixels

Procedure label-components(target component pixels):

1. X=[1,1,0,-1,-1,-1,0,1]
2. Y=[0,-1,-1,-1,0,1,1,1]
3. label_number = 2
4. Let L = rank target component pixels locations from min to max
5. For all pixel in L:
6. If pixel is not visited:
7. Let Q be a queue
8. Pixel.visit = True
9. Pixel.label = label_number
10. Q.enqueue(Pixel)
11. While Q is not empty:
12. Pixel = Q.dequeue()
13. For all x in X:
14. For all y in Y:
15. If pixels[i+x,j+y] is not visited:
16. Q.enqueue(pixels[i+x,j+y])
17. pixel[i+x,j+y].visit = True
18. pixel[i+x,j+y].label = label_number
19. label_number+=1

Tucker, Compton J. 1978. “Red and Photographic Infrared Linear Combinations for Monitoring Vegetation.”

Wan, Liang, Yijian Li, Haiyan Cen, Jiangpeng Zhu, Wenxin Yin, Weikang Wu, Hongyan Zhu, Dawei Sun, Weijun Zhou, and Yong He. 2018. “Combining UAV-Based Vegetation Indices and Image Classification to Estimate Flower Number in Oilseed Rape.” *Remote Sensing* 10 (9): 1484.

Woebbecke, David M, George E Meyer, K von Bargen, and D A Mortensen. 1995. “Color Indices for Weed Identification under Various Soil, Residue, and Lighting Conditions.” *Transactions of the ASAE* 38 (1): 259–69.
